# Live-cell fluorescence lifetime multiplexing using synthetic fluorescent probes

**DOI:** 10.1101/2022.01.15.476486

**Authors:** Michelle S. Frei, Birgit Koch, Julien Hiblot, Kai Johnsson

## Abstract

Fluorescence lifetime multiplexing requires fluorescent probes with distinct fluorescence lifetimes but similar spectral properties. Even though synthetic probes for many cellular targets are available for multicolor live-cell fluorescence microscopy, few of them have been characterized for their use in fluorescence lifetime multiplexing. Here we demonstrate that from a panel of 18 synthetic probes, eight pairwise combinations are suitable for fluorescence lifetime multiplexing in living mammalian cell lines. Moreover, combining multiple pairs in different spectral channels enables us to image up to six different biological targets, effectively doubling the number of observable targets. The combination of synthetic probes with fluorescence lifetime multiplexing is thus a powerful approach for live-cell imaging.

## Introduction

Fluorescence microscopy is an indispensable tool to non-invasively investigate dynamic processes in living cells. Such experiments often require to image multiple biomolecules and cellular compartments simultaneously. This is generally achieved by spectrally resolved detection using fluorophores with distinct excitation and emission spectra (Figure 1A). However, even though fluorophores that cover the entire visible spectrum are available^1,2^, this approach is often limited to three to four channels as the spectra of the fluorophores overlap^3^. Strategies to expand the degree of multiplexing have not only centered on techniques to improve spectral imaging^4–6^, but also make use of other fluorophore properties to access higher dimensions. One such property is fluorescence lifetime, which has been used for multiplexing via fluorescence lifetime imaging microscopy (FLIM, Figure 1B)^7,8^. Recently, lifetime multiplexing was combined with spectral multiplexing (S-FLIM) to further increase the number of simultaneously observable targets^9,10^.

**Figure 1.**
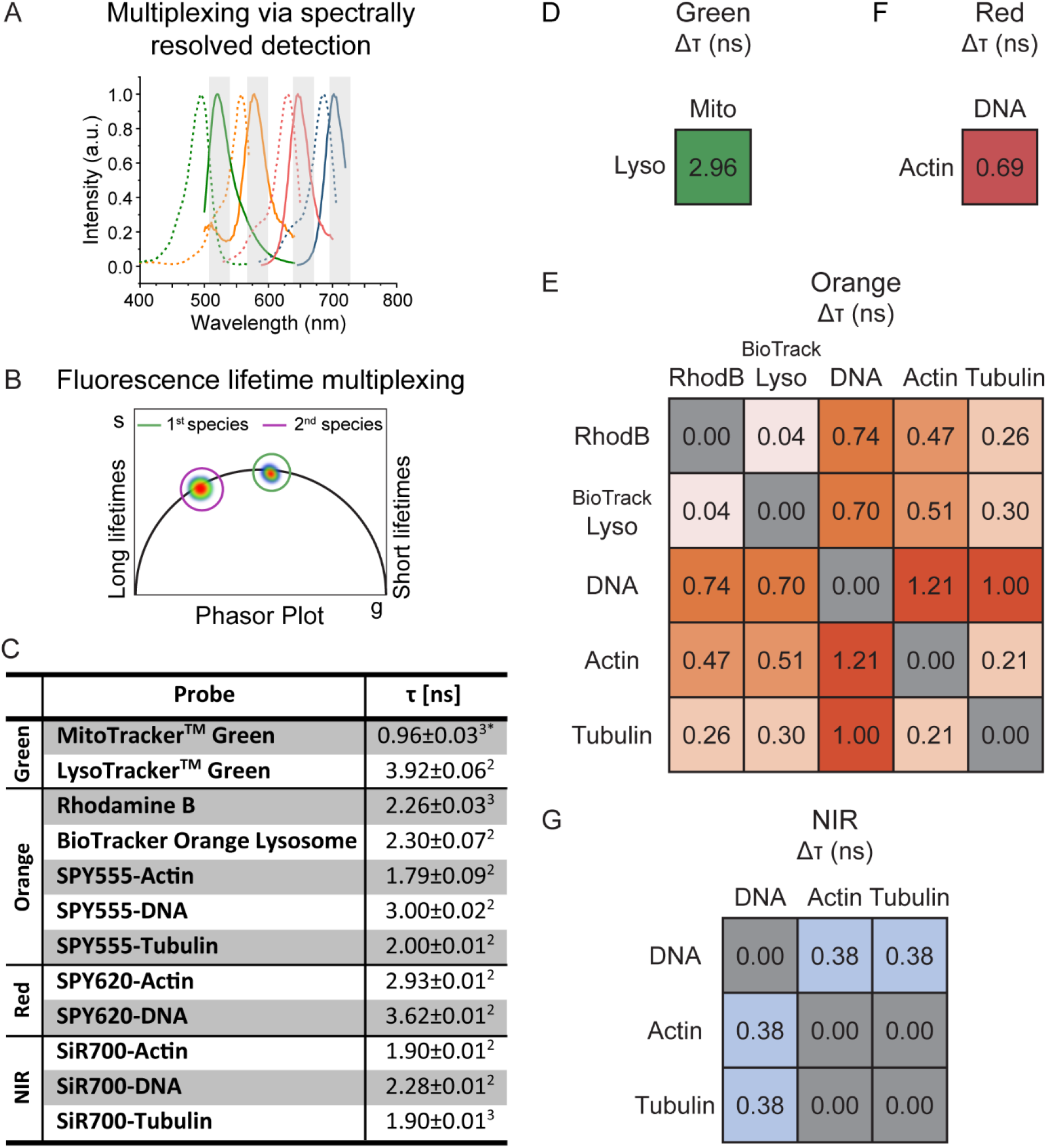
Fluorescence lifetime characterization of synthetic probes. **A**-**B** Schematic representation of multiplexing via spectrally resolved detection (**A**) or fluorescence lifetime multiplexing (**B**). **C** Average intensity weighted fluorescence lifetime (τ) of the 12 probes suitable for fluorescence lifetime multiplexing. Mean±s.e.m., *N* = 4 field of views from 2 biological replicates.^2^ bi-exponential fit, ^3^ tri-exponential fit, * tail fit (all others n-exponential reconvolution fit). **D-G** Differences in average intensity weighted fluorescence lifetime (Δτ) between probes in the green (**D**), orange (**E**), red (**F**), and NIR (**D**) spectral region.

Synthetic probes based on small-molecule fluorophores for live-cell microscopy of various subcellular targets such as lysosomes, mitochondria, or filamentous actin (F-actin) are available^11^. They do not require genetic engineering (e.g. transfection) of the target cell and can therefore be applied to a wide variety of cell types. Additionally, probes for different targets can easily be combined, while the simultaneous expression of multiple tagged proteins can be challenging^12^. Indeed, synthetic probes with distinct spectral properties were successfully combined for multiplexing^13–15^. However, they have only found limited use in fluorescence lifetime multiplexing and their fluorescence lifetimes are often not characterized^9,10,16,17^. Proof-of-concept studies were restricted to fixed cell applications^9,10^ and/or used probes with both differences in fluorescence lifetime and emission spectrum^9,10,16^. We recently demonstrated the combined use of synthetic probes and self-labeling protein tags for fluorescence lifetime multiplexing.^17^ However, our study was limited to only four probes and thus only partially exploited the potential of synthetic probes for fluorescence lifetime multiplexing.

Here, we investigate if fluorophores of different classes and chemically identical fluorophores targeted to different subcellular localizations show differences in fluorescence lifetime. Indeed, internal and external factors including vibrational and rotational freedom, viscosity, polarity, or the presence of quenching moieties can influence the fluorescence lifetime of fluorophores^18^. Combinations of live-cell compatible, synthetic probes for biomolecules or cellular compartments could find applications in live-cell fluorescence lifetime multiplexing. Ideally, each probe should show a homogenous and narrow fluorescence lifetime distribution and spectrally similar probes should show differences in fluorescence lifetimes to enable their separation.

## Results

We therefore characterized the spectral and fluorescence lifetime properties of 18 commercially available cell permeable probes. These encompassed popular rhodamine and BODIPY based probes in five different spectral channels targeting DNA, F-actin, microtubules, mitochondria, and lysosomes (Supporting Table S1)^13,15,19,20^. Four probes were previously used for fluorescence lifetime multiplexing^16,17^ and another two were used in spectral FLIM (S-FLIM)^9,10^. For initial screening purposes we choose to assess the fluorescence lifetime properties by phasor analysis^21,22^ as it allows to rapidly and visually screen probes for differences in fluorescence lifetime without the need for fitting (Figure 1B, Supporting Figure S1). Specifically, we labeled living U-2 OS cells with all 18 probes individually and acquired FLIM images. Phasor plot analysis then revealed narrow and homogenous distributions for all actin, microtubule, and DNA probes. Probes for lysosomes had slightly broader distributions but were still homogeneous. However, only two of the four probes for mitochondria displayed satisfying properties (MitoTracker^TM^-Green and rhodamine B). The other two, MitoTracker^TM^-Red and MitoTracker^TM^-Orange, showed multiple fluorescence lifetime populations in their phasor plots (Supporting Figure S1). This might result from the probes accumulating in multiple organelles as previously reported for MitoTracker^TM^-Orange in fixed cells (mitochondria, nucleoli, and endosome)^9^. While these differences in fluorescence lifetime might be used to separate the specific mitochondria signal from the unspecific signal in the endosome and the nucleoli, these probes cannot be combined with other spectrally similar probes for further fluorescence lifetime multiplexing and hence they were not further investigated.

Next, we assessed the differences in fluorescence lifetime within one spectral channel by overlaying the measured phasor plots of the pure species. If differences were found, the individual average intensity-weighted fluorescence lifetime was quantified by curve fitting. This then allowed to calculate the fluorescence lifetime differences between probes in the same spectral channel. The largest differences in fluorescence lifetime were found in the green spectral channel between LysoTracker^TM^-Green (3.92±0.06 ns) and MitoTracker^TM^-Green (0.96±0.03 ns) followed by SPY555-DNA (3.00±0.02 ns) and SPY555-Actin (1.79±0.09 ns) in the orange channel. These two pairs are hence ideally suited for fluorescence lifetime multiplexing (Figure 1C-E). Differences were also found for the SPY620 and SiR700 probes (Figure 1F-G). Probes based on the popular silicon rhodamine (SiR) did not show any differences in fluorescence lifetime and were therefore not further investigated (Supporting Figure S1). Generally, probes based on different fluorophore scaffolds (e.g. BODIPY vs. rhodamine) showed the biggest differences in fluorescence lifetime. The excitation and emission spectra of probes within one spectral region were generally highly similar, except for LysoTracker^TM^-Red, which shows a 20 nm hypsochromic shift in comparison to the two SPY620 probes (Supporting Figure S2). LysoTracker ^TM^-Red was therefore not considered for multiplexing experiments. Furthermore, we demonstrated that most probes’ fluorescence lifetime showed only little variation between different cell lines (HeLa and HEK293, Supporting Table S2). An exception are the lysosome probes, which generally showed broad lifetime distributions (Supporting Figure S1) and therefore higher variability in average intensity weighted fluorescence lifetime within one cell line as well as between different cell lines. This variability might stem from differences in the intralysosomal pH^23^.

We then performed fluorescence lifetime multiplexing using pairwise combinations of two probes in each of the four spectral regions (green: 489 nm excitation, 510-540 nm emission; orange: 550 nm excitation, 570-600 nm emission; red: 615 nm excitation, 635-700 nm emission; near-infrared (NIR): 670 nm excitation, 710-760 nm emission). As predicted by the differences in average intensity weighted fluorescence lifetime (Figure 1D-G), multiple combinations of probes could be separated using fluorescence lifetime information. Suitable pairs were found in all four spectral regions (Figure 2A-D). For instance, it was possible to image mitochondria and lysosomes simultaneously using LysoTracker^TM^-Green and MitoTracker^TM^-Green in the green channel (Figure 2A). The nucleus and F-actin can be separated using probes in either the orange, the red, or the NIR channel (Figure 2C, Supporting Figure S3-4). Larger differences in fluorescence lifetime facilitate separation but the brightness of the different species also plays a role. In order to obtain comparable signal-to-noise ratios for both species after separation, similar photon numbers should be collected. This can be challenging when working with probes of different brightness as the fluorescence of both probes is acquired simultaneously using only one excitation wavelength. However, the *in cellulo* brightness of synthetic probes is not only determined by their molecular brightness but can also be adjusted through the degree/density of labeling and hence the labeling concentration. Specifically, mitochondria and lysosome probes accumulate in their respective organelle. The labeling concentration and hence the brightness can therefore be varied over a broad range. The brightness of DNA, actin, and tubulin probes, on the other hand, cannot be varied to the same degree as their number of binding sites is limited. Moreover, most probe combinations showed large enough differences in fluorescence lifetime (>0.5 ns) such that the two species can be directly distinguished using average photon arrival times as reported in the FastFLIM image. This allows to distinguish the two species during image acquisition without the need for fitting or phasor analysis, greatly simplifying the use of fluorescence lifetime multiplexing. As the probes’ fluorescence lifetimes showed little variation between cell types (Supporting Table S2), multiplexing could also be performed in living HEK 293 or HeLa cells (Supporting Figure S5).

**Figure 2.**
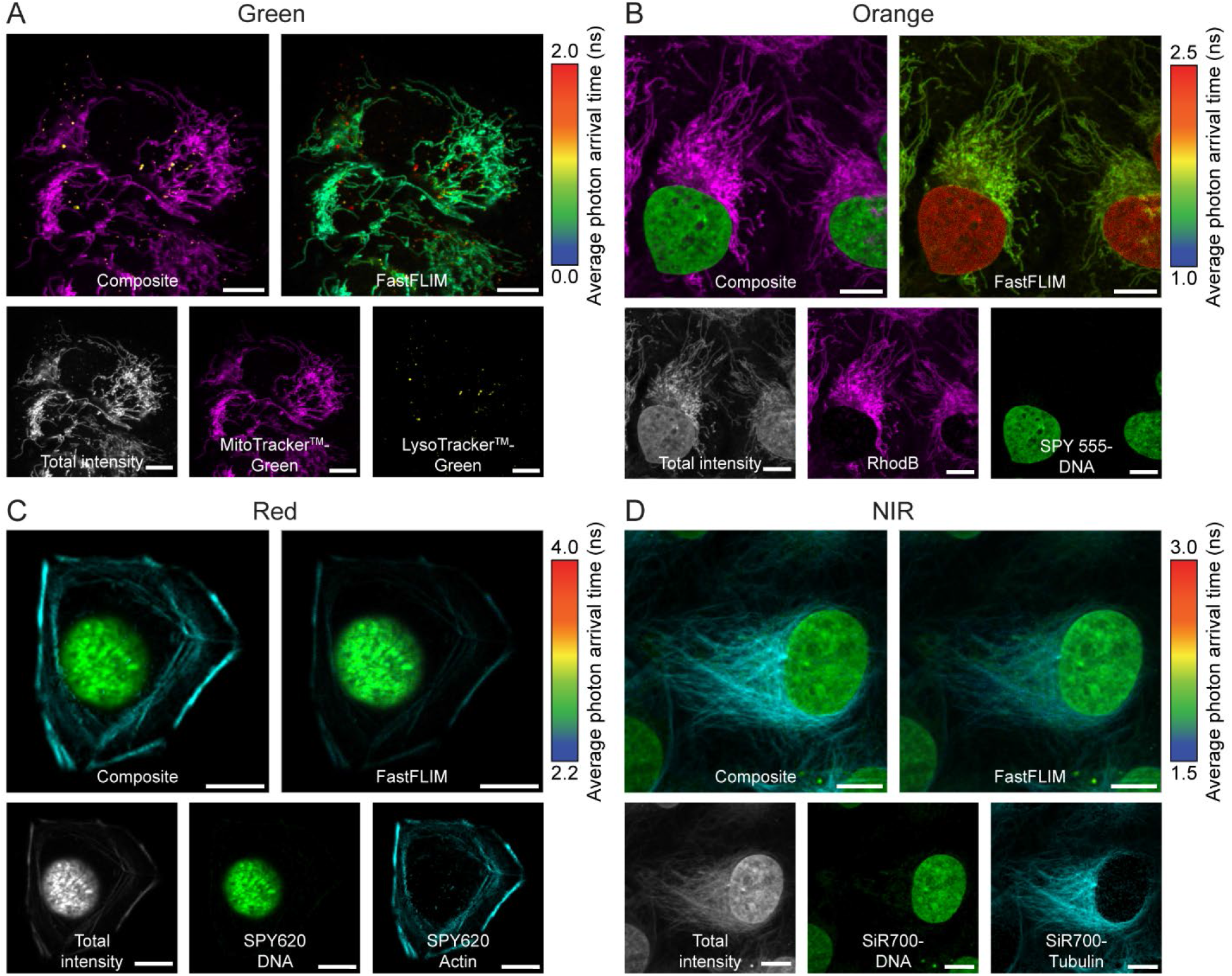
Live-cell fluorescence lifetime multiplexing in four different spectral regions. **A-D** U-2 OS cells were labeled with MitoTracker^TM^-Green and LysoTracker^TM^-Green (**A**), Rhodamine B and SPY555-DNA (**B**), SPY620-DNA and SPY620-Actin (**C**), or SiR700-DNA and SiR700-Tubulin (**D**) and imaged by FLIM. In each spectral channel the two species could be clearly separated based on fluorescence lifetime information. The FastFLIM image reports the average photon arrival time and allows for quick visual inspection of the two species. The composite, the FastFLIM image with the respective color-scale, the total fluorescence intensity, and the two individual separated species are given. Species separation was achieved using the phasor approach (positioning the cluster-circles on the phasor plot at the position of the pure species). Scale bars, 10 μm.

The herein tested probes allowed to simultaneously image combinations of mitochondria, lysosomes, nucleus (DNA), F-actin, or microtubules. In order to access alternative targets for which no synthetic probes are available or for which the probes do not show a difference in fluorescence lifetime, the self-labeling protein (SLP) tag strategy can be employed. SLP tags, such as HaloTag7^24^ or SNAP-tag^25^, react with fluorophores bearing a chloroalkane (CA) or benzylguanine (BG) ligand, respectively. Through fusion of the SLP tag to proteins of interest (POI) one can therefore localize cell-permeable fluorophores to different subcellular localizations. We previously demonstrated the use of SLP-tags in fluorescence lifetime multiplexing and characterized the average fluorescence lifetime of different HaloTag7 fusion proteins conjugated to MaP555-CA (2.4 ns) or MaP618-CA (3.1 ns)^17^. We hence performed a similar characterization for SNAP-tag localized to different subcellular localizations and labeled with MaP555-BG in living U-2 OS cells (e.g. histone 2B, Lamin B1, Tomm20 etc.; Supporting Table S3). SNAP-tag-MaP555 showed little variation in fluorescence lifetime averaging around 2.5 ns. The MaP555 substrates for SNAP (BG) and HaloTag (CA) should therefore be multiplexable with the SPY555 probes (Actin, DNA, and Tubulin) as they have a fluorescence lifetime difference of around 0.5 ns. To test this, we expressed HaloTag7-SNAP-tag as a fusion with the endoplasmic reticulum (ER) marker calreticulin (CalR)/KDEL in U-2 OS cells and labeled them with either MaP555-BG or MaP555-CA. When combined with SPY555-Actin or SPY555-DNA, two species could be separated based on their fluorescence lifetime information (Supporting Figure S6). On the other hand, the fluorescence lifetimes of Rhodamine B and BioTracker Orange Lysosome are too similar to the corresponding SLP probes and therefore did not allow multiplexing. MaP618-CA (3.1 ns) can be used for multiplexing with SPY620-DNA but not SPY620-Actin (Supporting Figure S7).

We then combined fluorescence lifetime multiplexing with spectrally resolved detection. First, four probes were multiplexed in two spectral channels each containing two probes separable in fluorescence lifetime (Figure 3A): LysoTracker^TM^-Green and MitoTracker^TM^-Green with two SPY555 probes (e.g. SPY555-DNA and SPY555-Tubulin). Living U-2 OS cells labeled with this combination were imaged in both the green and orange channel by FLIM. Separation of the two lifetime components in each spectral channel gave access to a four species image of mitochondria, lysosomes, the nucleus, and the microtubule network (Supporting Figure S8). Alternatively, SPY555-Tubulin could be replaced by SPY555-Actin revealing the F-actin network. Instead of combining the green and the orange channel, two red probes (SPY620-DNA and SPY620-Actin) can also be combined with the two probes in the green channel (LysoTracker^TM^-Green and MitoTracker^TM^-Green; Supporting Figure S9). The expression of HaloTag7-SNAP-tag in the ER of U-2 OS cells and labeling with LysoTracker^TM^-Green, MitoTracker^TM^-Green, MaP555-CA, SPY555-Actin, SiR700-DNA, and SiR700-Tubulin even allowed to acquire six species images. Acquisition of all three channels and separation of the two lifetime components in each of them revealed lysosomes, mitochondria, the ER, F-actin, the nucleus, and microtubules (Figure 3B, Supporting Figure S10). This strategy hence allows to double the number of species imaged in spectrally distinct channels.

**Figure 3.**
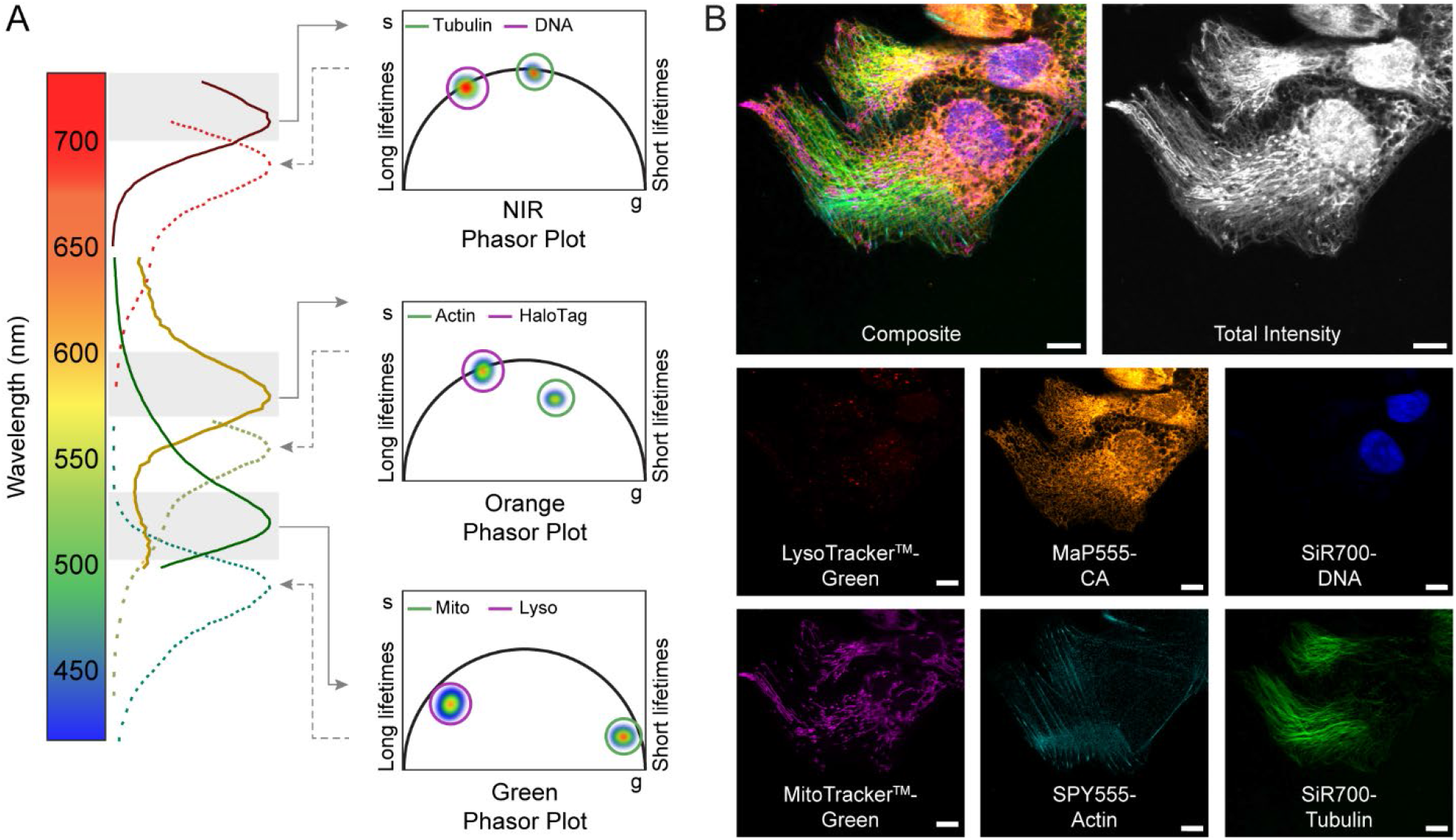
Combination of spectrally resolved detection and fluorescence lifetime multiplexing. **A** Schematic view of fluorescence lifetime multiplexing combined with spectrally resolved detection in three channels. **B** Fluorescence lifetime multiplexing of U-2 OS cells stably expressing a HaloTag7-SNAP-tag fusion in the ER. Cells were labeled with LysoTracker^TM^-Green, MitoTracker^TM^-Green, MaP555-CA, SPY555-Actin, SiR700-DNA, and SiR700-Tubulin. The composite, the total fluorescence intensity, and the six individual separated species are given. Species separation was achieved using the phasor approach (positioning the cluster-circles on the phasor plot at the position of the pure species). Scale bars, 10 μm.

In summary, the fluorescence lifetimes of 18 synthetic fluorescent probes were characterized and eight pairwise combinations of probes suitable for fluorescence lifetime multiplexing identified. In combination with spectrally resolved detection, these synthetic probes allow to double the number of species that can be imaged giving access to two, four, and even six species images. As more and more probes based on a variety of different fluorophores become available, we believe that it should be possible to expand this number further. The use of synthetic probes targeting different biomolecules or cellular compartments is hence a straightforward strategy to generate fluorescence lifetime contrast and should facilitate the use of fluorescence lifetime multiplexing in living cells.

## Methods

### General considerations

MaP555-BG, MaP555-CA, and MaP618-CA were prepared according to literature procedures^15^ by B. Réssy or D. Schmidt (MPI-MRI). All other probes were purchased from commercial vendors (Supporting Table S1), or obtained from Spirochrome. Fluorophores were prepared as stock solutions in dry DMSO and diluted in imaging medium such that the final concentration of DMSO did not exceed 1% v/v.

### Plasmids

A pcDNA5/FRT/TO vector (ThermoFisher Scientific) was used for transient expression and a pcDNA5/FRT vector (ThermoFisher Scientific) for stable cell line establishment in mammalian cells. HaloTag7-Pro30-SNAP-tag was fused to CalR/KDEL for ER localization. SNAPf-tag was fused to CEP41, H2B, TOMM20, NES, COX8, CalR/KDEL, β4Gal-T1, LAMP1, SKL, Lyn11, Ig-κ-/PDGFR, or Lifeact for expression in mammalian cells. Cloning was performed by Gibson assembly^26^. DNA was subsequently electroporated in *E. cloni* 10G (Lucigene) and plated on agar Lysogenic Broth plates with 100 μg/mL ampicillin and incubated at 37 °C overnight. CEP41 (Addgene plasmid # 135446)^27^, TOMM20 (Addgene plasmid # 135443)^27^, H2B (Addgene plasmid # 135444)^27^, COX8 (Addgene plasmid # 113916)^28^, Ig-κ-/-PDGFR^29^, and SNAPf (Addgene plasmid # 167271)^30^ were available in house and used as template plasmids. mCherry-LaminB1-10 was a gift from Michael Davidson (Addgene plasmid # 55069), pAAV_hsyn_NES-his-CAMPARI2-F391W-WPRE-SV40 was a gift from Eric Schreiter (Addgene plasmid # 101061)^31^, pmTurquoise2-Golgi was a gift from Dorus Gadella (Addgene plasmid # 36 2 05)^32^, and LAMP1-mGFP was a gift from Esteban Dell’Angelica (Addgene plasmid # 34831)^33^. For more information see Supporting Table S4. Plasmids generated in this work will be deposited on Addgene and the accession codes are listed in Supporting Table S4.

### Cell culture and transfection

U-2 OS (ATCC), HeLa (ATCC), HEK 293 (ATCC), and U-2 OS Flp-In TREx CalR-HaloTag7-Pro30-SNAP-tag-KDEL cells were cultured in high-glucose phenol red free DMEM (Life Technologies) medium supplemented with GlutaMAX (Life Technologies), sodium pyruvate (Life Technologies) and 10% FBS (Life Technologies) in a humidified 5% CO2 incubator at 37 °C. Cells were split every 3–4 days or at confluency. Cell lines were regularly tested for mycoplasma contamination. Cells were seeded on 8 well glass bottom dishes (Ibidi) or 96 well plates (Eppendorf) three to one day before imaging. Transient transfections were performed using Lipofectamine^TM^ 2000 reagent (Life Technologies) according to the manufacturer’s recommendations: the DNA (0.3 μg) was mixed with OptiMEM I (10 μL, Life Technologies) and Lipofectamine^TM^ 2000 (0.75 μL) was mixed with OptiMEM I (10 μL, 8 well). The solutions were incubated for 5 min at room temperature, then mixed and incubated for an additional 20 min at room temperature. The prepared DNA-Lipofectamine complex was added to one of the wells in an 8 well glass bottom dish with cells at 50–70% confluency. After 12 h incubation in a humidified 5% CO2 incubator at 37 °C the medium was changed to fresh medium. The cells were incubated under the same conditions for 24–48 h before imaging.

### Stable cell line establishment

The Flp-In^TM^ T-REx^TM^ System (ThermoFisher Scientific) was used to generate a stable cell line exhibiting expression of CalR-HaloTag7-Pro30-SNAP-tag-KDEL. Briefly, pcDNA5-FRT-GOI and pOG44 were co-transfected into the host cell line U-2 OS FlpIn TREx^34^. Homologous recombination between the FRT sites in pcDNA5-FRT-GOI and the host cell chromosome, catalyzed by the Flp recombinase expressed from pOG44, produced the U-2 OS FlpIn TREx cells expressing stable GOI. Selection was performed using 100 μg mL^-1^ hygromycin B (ThermoFisher Scientific) and 15 μg mL^-1^ blasticidine (ThermoFisher Scientific). The stable cell line was seeded on glass bottom dishes as described before.

### Labeling and sample preparation

Cells were labeled with the respective fluorophores:

- MitoTracker^TM^-Green: 200 nM, 30 min, 37 °C, one wash
- LysoTracker^TM^-Green: 75 nM, 30 min, 37 °C, one wash
- SPY555-Actin: 500 nM, 1 h, 37 °C, no wash
- SPY555-DNA: 1 μM, 1 h, 37 °C, no wash
- SPY555-Tubulin: 500 nM, 1 h, 37 °C, no wash
- MaP555-CA: 500 nM, 1 h, 37 °C, one wash
- MaP555-BG: 2 μM, 1 h, 37 °C, two washes
- MitoTracker^TM^-Orange: 500 nM, 30 min, 37 °C, one wash
- BioTracker 560 Orange Lysosomes: 10 μg of the fluorophore were diluted in 200 μL DMSO and a 1:1,000 dilution was used for labeling, 30 min, one wash
- Rodamine B: 1 μM, 30 min, 37 °C, one wash
- MitoTracker^TM^-Red: 500 nM, 30 min, 37 °C, one wash
- LysoTracker^TM^-Red: 75 nM, 30 min, 37 °C, one wash
- SPY620-Actin: 1 μM, 1 h, 37 °C, no wash
- SPY620-DNA: 500 nM, 1 h, 37 °C, no wash
- MaP618-CA: 500 nM, 1 h, 37 °C, one wash
- SiR-DNA: 500 nM, 1 h, 37 °C, no wash
- SiR-Tubulin: 500 nM, 1 h, 37 °C, no wash
- SiR-Actin: 500 nM, 1 h, 37 °C, no wash
- SiR700-DNA: 1 μM, 1 h, 37 °C, no wash
- SiR700-Tubulin: 500 nM, 1 h, 37 °C, no wash
- SiR700-Actin: 500 nM, 1 h, 37 °C, no wash

in phenol-red free DMEM medium supplemented with GlutaMAX, sodium pyruvate, and 10% FBS (all Life Technologies), washed with the same medium as indicated above. If no-wash and wash probes were used simultaneously, labeling with wash probes was performed first followed by labeling with no-wash probes. Imaging was performed in the same medium.

### Confocal microscopy

Confocal fluorescence microscopy was performed on a Leica SP8 FALCON microscope (Leica Microsystems) equipped with a Leica TCS SP8 X scanhead; a SuperK white light laser, Leica HyD SMD detectors, a HC PL APO CS2 40×1.10 water objective. Emission was collected as indicated in Supporting Table S5. The microscope was equipped with a CO2 and temperature controllable incubator (Life Imaging Services, 37 °C).

### Fluorescence excitation and emission spectra

Fluorescence excitation and emission spectra of synthetic probes were measured in living U-2 OS cells by confocal microscopy. For SNAP-tag and HaloTag7 probes U-2 OS cells were transiently transfected with HaloTag7 or SNAP-tag (no localization marker). Acquisition settings were as follows:

Green excitation: excitation between 470-534 nm in 2 nm steps; collection at 555 – 600 nm. Green emission: excitation at 470 nm and collection between 480–567 nm in 3 nm steps with a bandwidth of 10 nm.

Orange excitation: excitation between 475–575 nm in 2 nm steps; collection at 595–700 nm. Orange emission: excitation at 520 nm and collection between 530–617 nm in 3 nm steps with a bandwidth of 10 nm.

Red excitation: excitation between 550–650 nm in 2 nm steps; collection at 670–780 nm.

Red emission: excitation at 600 nm and collection between 610–697 nm in 3 nm steps with a bandwidth of 10 nm.

Except LysoTracker^TM^-Red excitation: excitation between 500–600 nm in 2 nm steps; collection at 620–700 nm. LysoTracker^TM^-Red emission: excitation at 540 nm and collection between 550– 647 nm in 3 nm steps with a bandwidth of 10 nm.

NIR excitation: excitation between 630–670 nm in 2 nm steps; collection at 720–780 nm.

NIR emission: excitation at 670 nm and collection between 690–768 nm in 3 nm steps with a bandwidth of 10 nm.

### Fluorescence lifetime imaging microscopy

FLIM was performed on a Leica SP8 FALCON microscope (as described above) at a pulse frequency of 80 MHz unless otherwise stated. Emission was collected as indicated in Supporting Table S5.

For determination of fluorescence lifetime cells were imaged, collecting 500 photons per pixel. The acquired images of cells were thresholded to remove background signal from empty coverslip space. Mean fluorescence lifetimes were calculated in the LAS X software (Leica Microsystems) by fitting a mono, bi, or tri-exponential decay model (n-exponential reconvolution, unless otherwise stated) to the decay (χ^2^ < 1.2).

For determination of average fluorescence lifetimes on different subcellular targets U-2 OS cells were transiently transfected with the SNAP-tag constructs and imaged, collecting 500 photons per pixel. The acquired images were processed as described above.

Structural images (species separation) were acquired as indicated in Supporting Table S5 and species separation was performed via phasor analysis positioning the cluster-circles on the phasor plot at the position of the pure species (Leica Microsystems)^21,22,35^.

### Software and image processing

All images were processed in LASX and ImageJ/Fiji^36,37^ unless otherwise stated. Excitation and emission spectra were visualized using OriginLab^38^.

## Data availability

Plasmids encoding SNAP-tag fusions will be deposited on Addgene. Correspondence and requests for the cell line should be addressed to K.J.

## Acknowledgement

This work was supported by the Max Planck Society. M.S.F was supported by the Deutsche Forschungsgemeinschaft (DFG, German Research Foundation) SFB TRR 186. The authors thank A. Bergner, B. Réssy, and D. Schmidt, for providing reagents.

## Contributions

M.S.F performed confocal and FLIM microscopy and the analysis thereof, B.K. generated the stable cell line, J.H. performed cloning of SNAP-tag fusions. M.S.F. and K.J. wrote the manuscript with input from all authors.

