## Supporting information for "Live-cell fluorescence lifetime multiplexing using synthetic fluorescent probes"

* Co-corresponding

### Table of Contents

##### Supporting Figures

**Figure S1:** Phasor plots of the 18 synthetic probes tested. S2

**Figure S2:** Spectral characterization of synthetic probes used for fluorescence lifetime multiplexing by confocal microscopy. S3

**Figure S3:** Live-cell fluorescence lifetime multiplexing using synthetic probes in different spectral regions. S4

**Figure S4:** Live-cell fluorescence lifetime multiplexing using synthetic probes in the orange spectral region. S5

**Figure S5:** Live-cell fluorescence lifetime multiplexing in different cell lines. S6

**Figure S6:** Multiplexing synthetic probes and self-labeling protein tags. S7

**Figure S7:** Multiplexing synthetic probes and HaloTag in the red channel. S8

**Figure S8:** Four species images combining fluorescence lifetime multiplexing in the green and orange spectral channel. S9

**Figure S9:** Four species images combining fluorescence lifetime multiplexing in the green and red spectral channel. S10

**Figure S10:** Six species images combining fluorescence lifetime multiplexing in the green, orange and NIR spectral channel. S11

##### Supporting Tables

**Table S1:** Structures of all 18 synthetic probes tested. S12-15

**Table S2:** Comparison of intensity weighted fluorescence lifetimes (τ) of different synthetic probes in three different cell lines. S16

**Table S3:** Comparison of intensity weighted fluorescence lifetimes (τ) of MaP555-BG on SNAPf-tag fused to different proteins of interest. S17

**Table S4:** Plasmids used and generated as well as the stable cell lines derived thereof. S18

**Table S5:** Fluorescence microscopy data acquisition parameters. S19-25

**References** S26

#### Supporting Figures

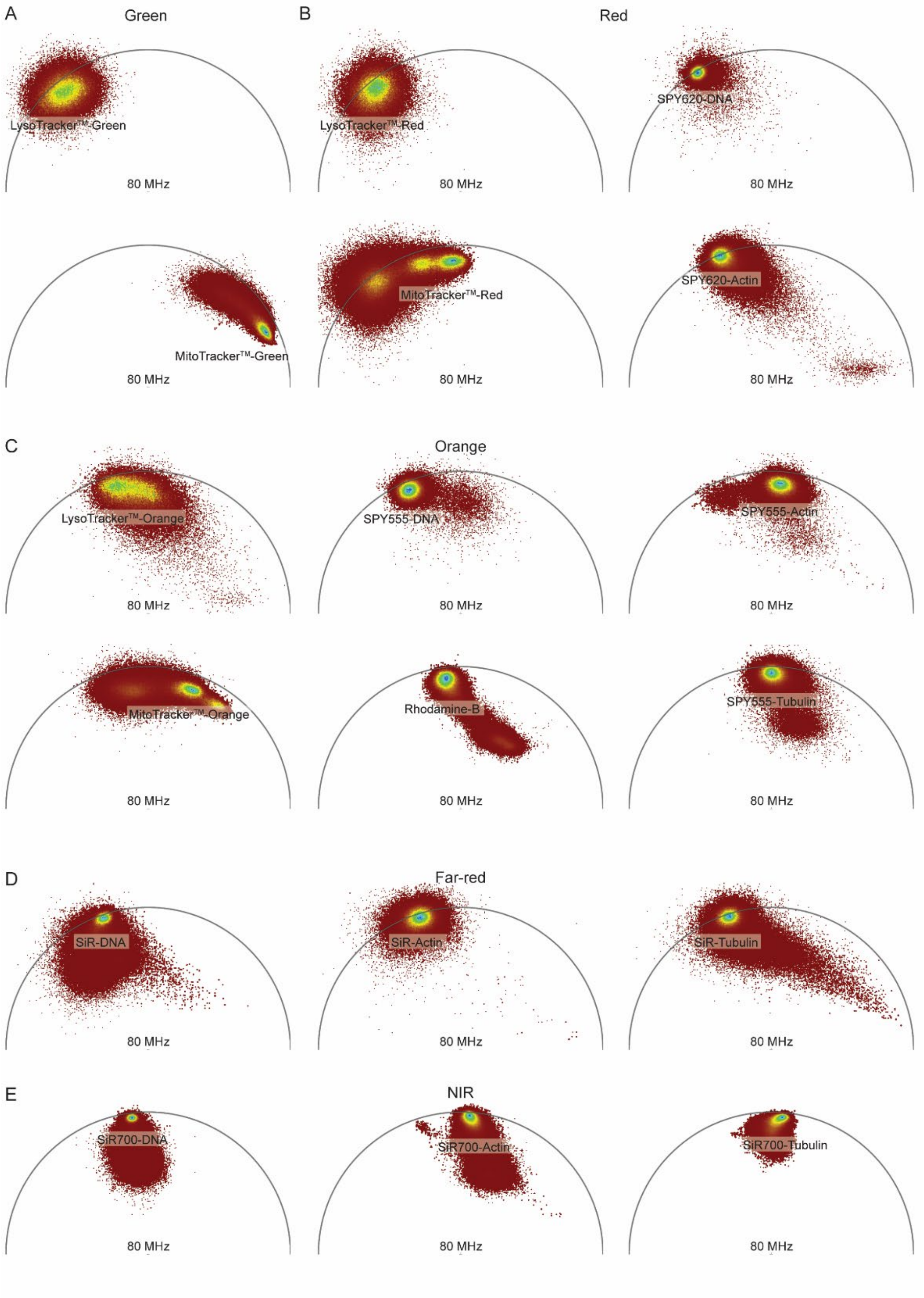

**Supporting Figure S1:** Phasor plots of the 18 synthetic probes tested. **A** Green spectral region. **B** Red spectral region. **C** Orange spectral region. **D** Far-red spectral region. **E** NIR spectral region.

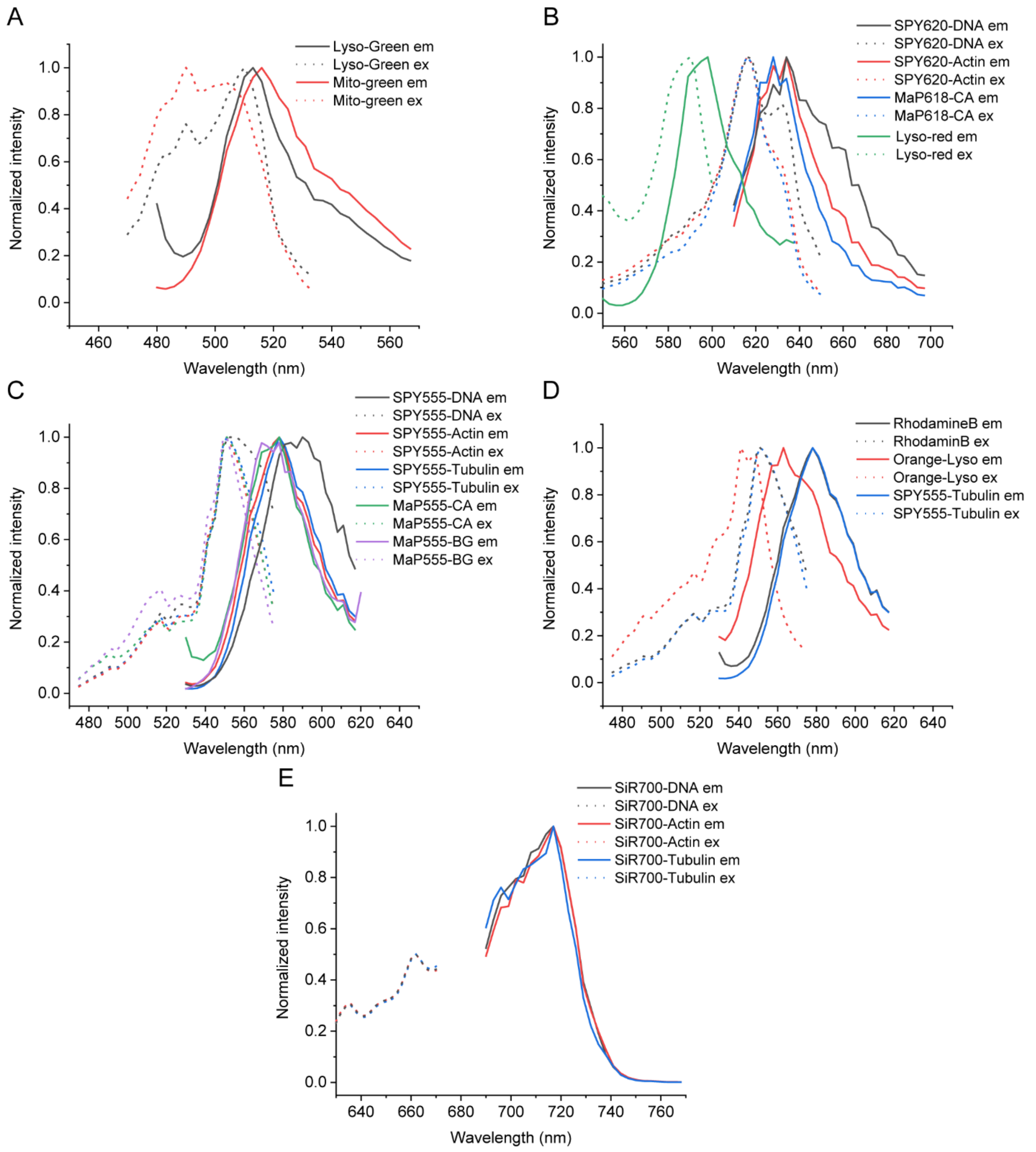

**Supporting Figure S2:** Spectral characterization of synthetic probes used for fluorescence lifetime multiplexing by confocal microscopy. **A-E** Normalized emission and excitation spectra of green probes (**A**), red probes (**B**), orange probes (**C**), orange probes (**D**), and SiR700 based NIR probes (**E**). For comparison of the orange probes the spectra of SPY555-Tubulin are given both in (**C**) and (**D**). The excitation maximum of the SiR700 based probes (~680 nm) could not be measured as the laser only reached 670 nm. The spectra were normalized to 0.5 instead of 1.0.

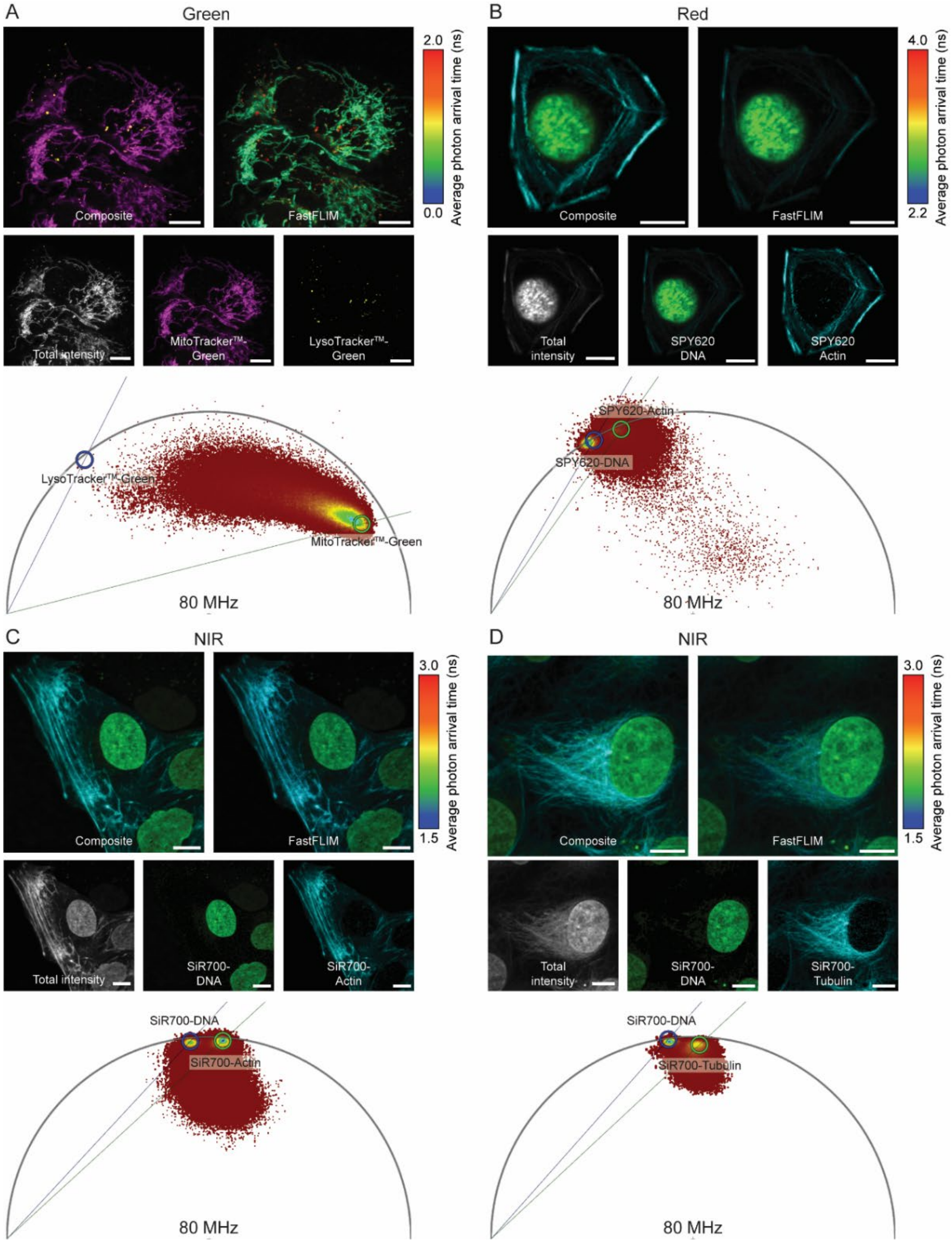

**Supporting Figure S3:** Live-cell fluorescence lifetime multiplexing using synthetic probes in different spectral regions. **A** Green spectral region labeling U-2 OS cells with MitoTracker$^{TM}$-Green and LysoTracker$^{TM}$-Green. **B** Red spectral region labeling U-2 OS cells with SPY620-DNA and SPY620-Actin. **C-D** NIR spectral region labeling U-2 OS cells with SiR700-DNA and SiR700-Actin (**C**) or SiR700-DNA and SiR700-Tubulin (**D**). The composite, the FastFLIM image with the respective color-scale, the total fluorescence intensity, the two individual separated species as well as the corresponding wavelet-filtered phasor plot used for separation are given. Species separation was achieved using the phasor approach by positioning the cluster-circles on the phasor plot at the position of the pure species. Scale bars, 10 µm.

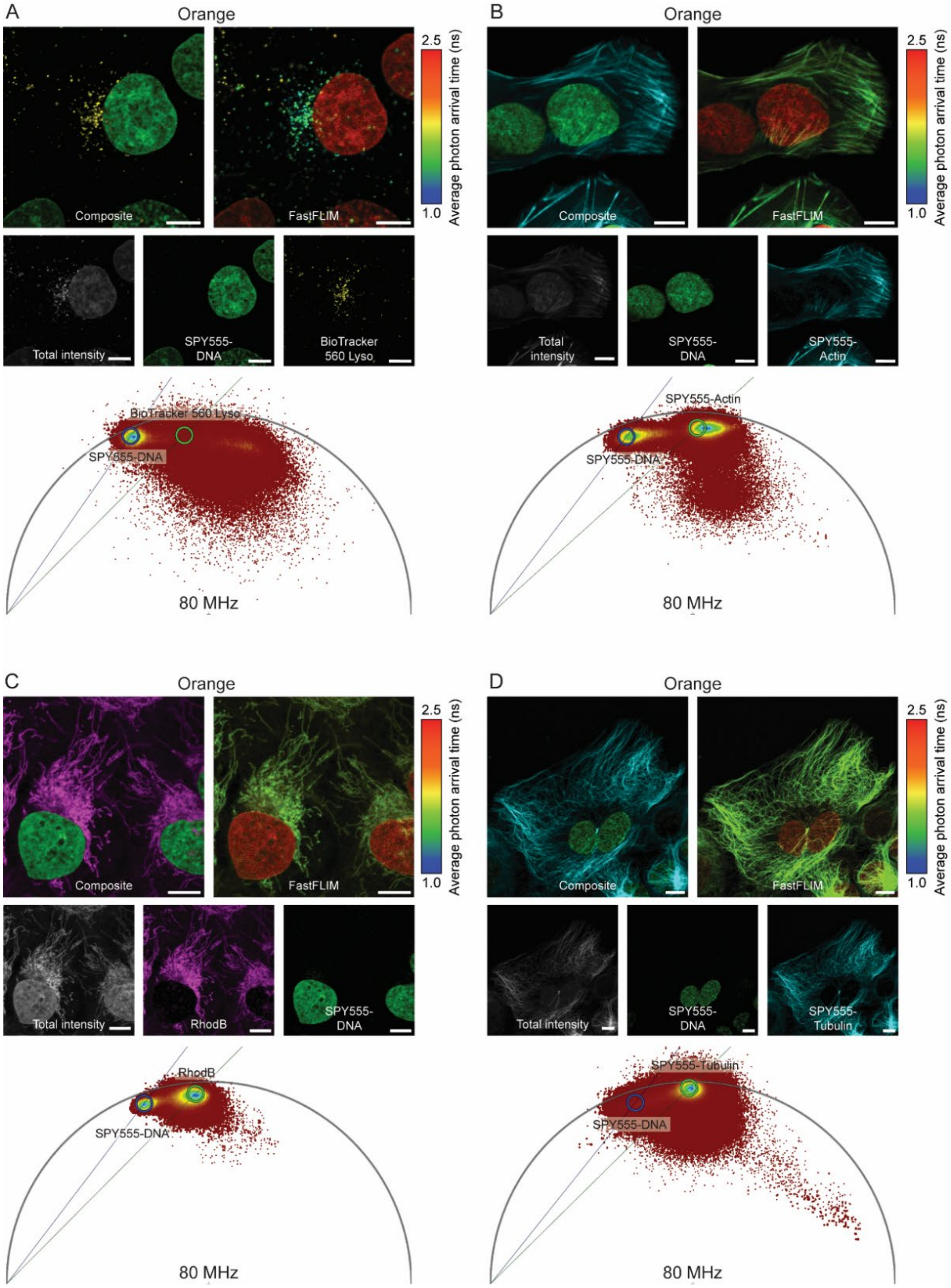

**Supporting Figure S4:** Live-cell fluorescence lifetime multiplexing using synthetic probes in the orange spectral region. **A-D** U-2 OS cells labeled with SPY555-DNA and BioTracker 560 Orange Lysosome (**A**), SPY555-DNA and SPY555-Actin (**B**), SPY555-DNA and RhodamineB (**C**), or SPY555-DNA and SPY55-Tubulin (**D**). The composite, the FastFLIM image with the respective color-scale, the total fluorescence intensity, the two individual separated species as well as the corresponding wavelet-filtered phasor plot used for separation are given. Species separation was achieved using the phasor approach by positioning the cluster-circles on the phasor plot at the position of the pure species. Scale bars, 10 µm.

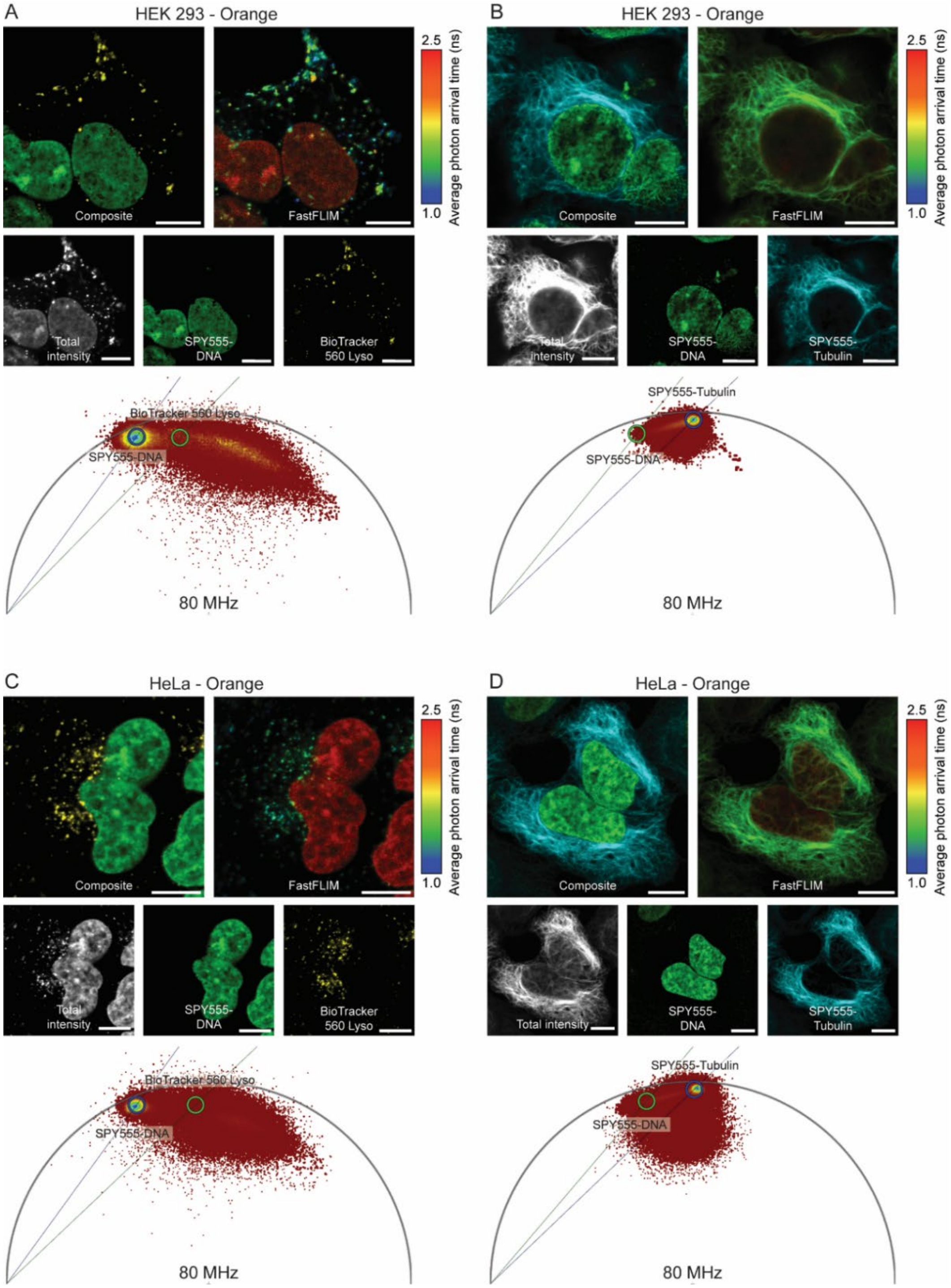

**Supporting Figure S5:** Live-cell fluorescence lifetime multiplexing in different cell lines. **A-D** HEK 293 (**A-B**) or HeLa cells (**C-D**) labeled with SPY555-DNA and BioTracker 560 Orange Lysosome (**A**, **C**) or SPY555-DNA and SPY555-Tubulin (**B**, **D**). The composite, the FastFLIM image with the respective color-scale, the total fluorescence intensity, the two individual separated species as well as the corresponding wavelet-filtered phasor plot used for separation are given. Species separation was achieved using the phasor approach by positioning the cluster-circles on the phasor plot at the position of the pure species. Scale bars, 10 µm.

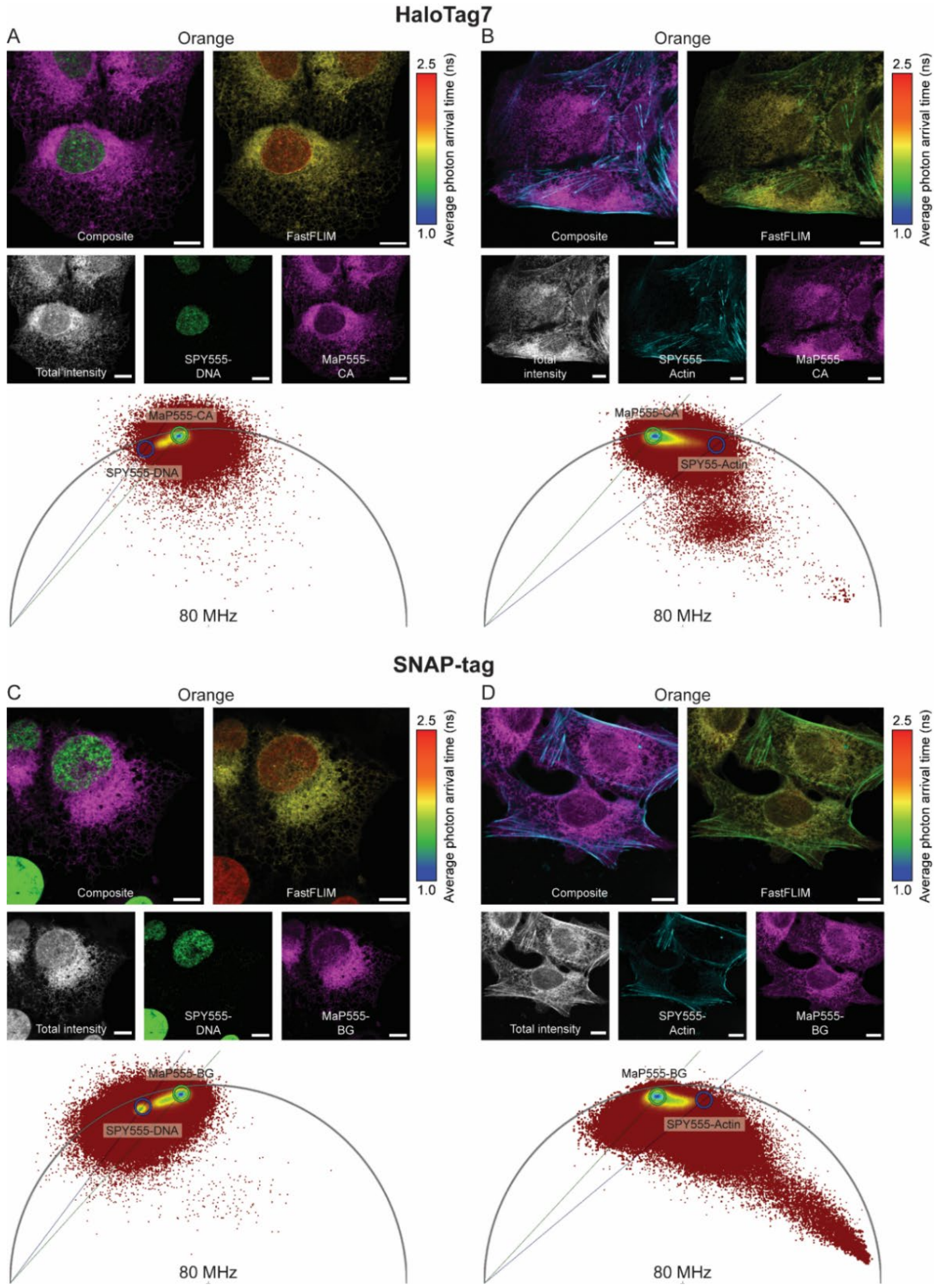

**Supporting Figure S6:** Multiplexing synthetic probes and self-labeling protein tags. **A-D** U-2 OS cells stably expressing the endoplasmic reticulum marker calreticulin (CalR) as a HaloTag7-SNAP-tag fusion additionally fused to a KDEL targeting peptide were labeled with MaP555-CA (**A**-**B**) or MaP555-BG (**C**-**D**) and SPY555-DNA (**A**, **C**) or SPY555-Actin (**B**, **D**). The composite, the FastFLIM image with the respective color-scale, the total fluorescence intensity, the two individual separated species as well as the corresponding wavelet-filtered phasor plot used for separation are given. Species separation was achieved using the phasor approach by positioning the cluster-circles on the phasor plot at the position of the pure species. Scale bars, 10 µm.

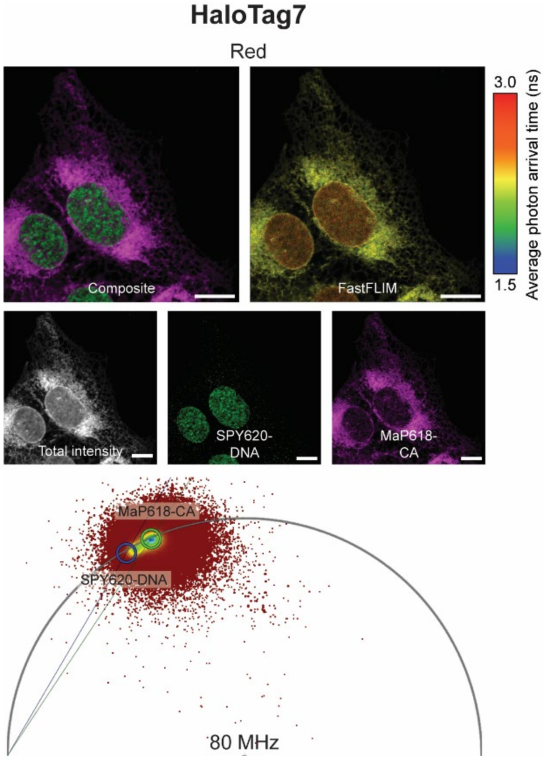

**Supporting Figure S7:** Multiplexing synthetic probes and HaloTag in the red channel. **A** U-2 OS cells stably expressing CalR-HaloTag7-SNAP-tag-KDEL were labeled with MaP618-CA and SPY620-DNA. The composite, the FastFLIM image with the respective color-scale, the total fluorescence intensity, the two individual separated species as well as the corresponding wavelet-filtered phasor plot used for separation are given. Species separation was achieved using the phasor approach by positioning the cluster-circles on the phasor plot at the position of the pure species. Scale bars, 10 µm.

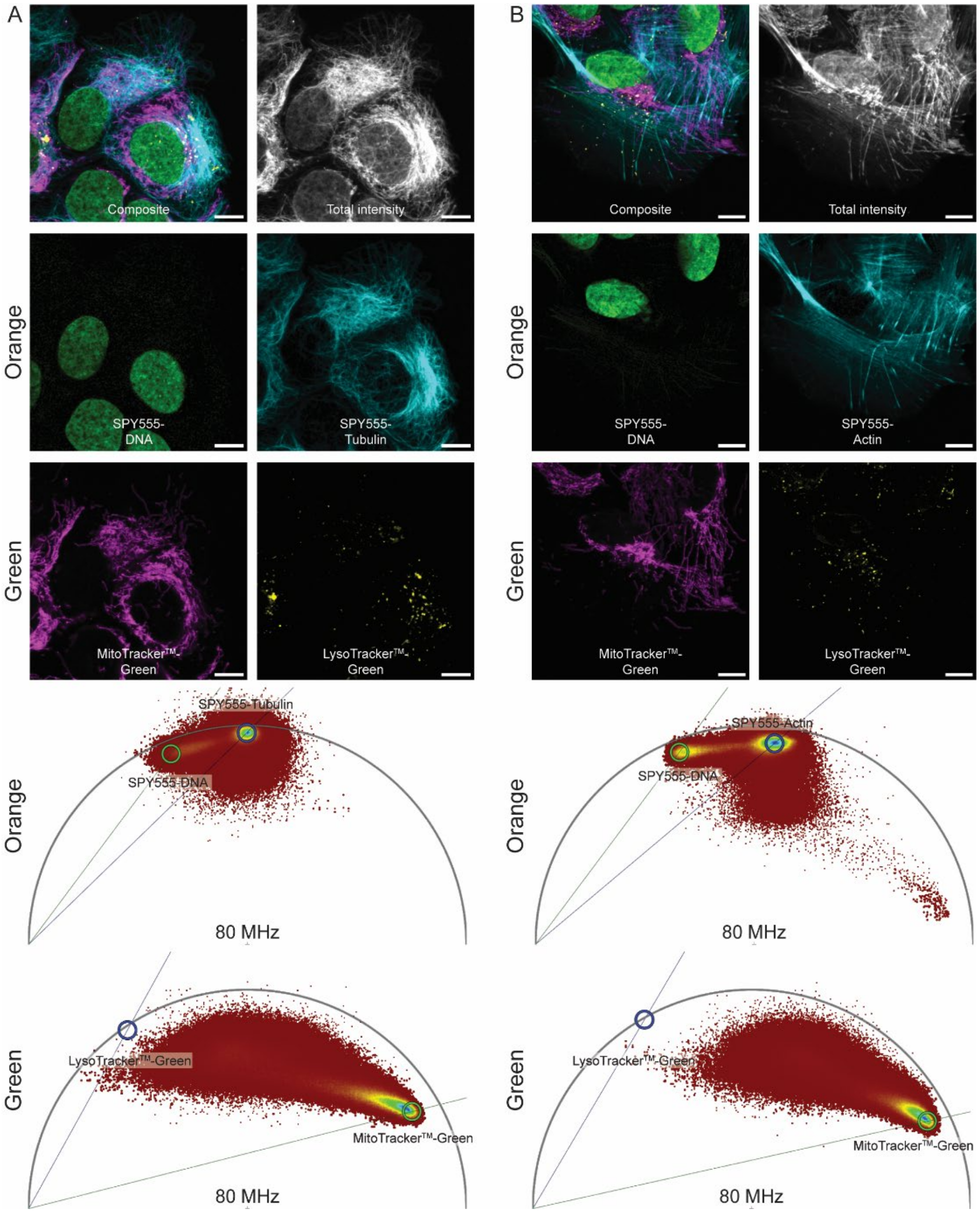

**Supporting Figure S8:** Four species images combining fluorescence lifetime multiplexing in the green and orange spectral channel. **A-B** U-2 OS cells were labeled with MitoTracker™-Green, LysoTracker™-Green, SPY555-DNA, and either SPY555-Tubulin (**A**) or SPY555-Actin (**B**) and imaged in the two spectral channels giving access to four species images. The composite, the total fluorescence intensity, the four individual separated species as well as the corresponding wavelet-filtered phasor plots used for separation are given. Species separation was achieved using the phasor approach by positioning the cluster-circles on the phasor plot at the position of the pure species. Scale bars, 10 µm.

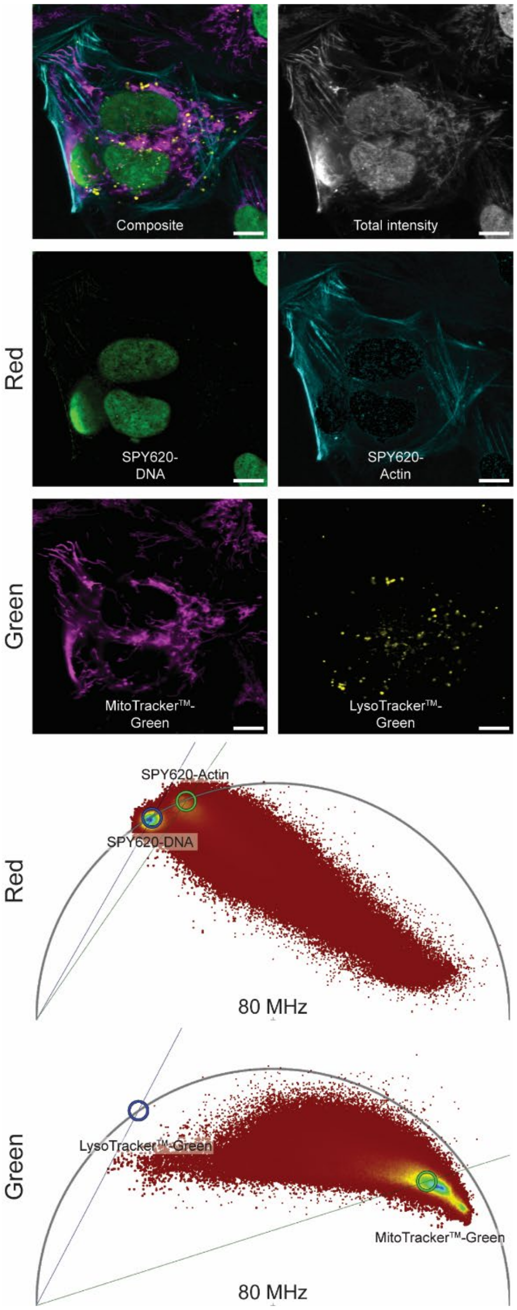

**Supporting Figure S9:** Four species images combining fluorescence lifetime multiplexing in the green and red spectral channel. U-2 OS cells were labeled with MitoTracker™-Green, LysoTracker™-Green, SPY620-DNA, and SPY620-Actin and imaged in the two spectral channels giving access to a four species image. The composite, the total fluorescence intensity, the four individual separated species as well as the corresponding wavelet-filtered phasor plots used for separation are given. Species separation was achieved using the phasor approach by positioning the cluster-circles on the phasor plot at the position of the pure species. Scale bars, 10 µm.

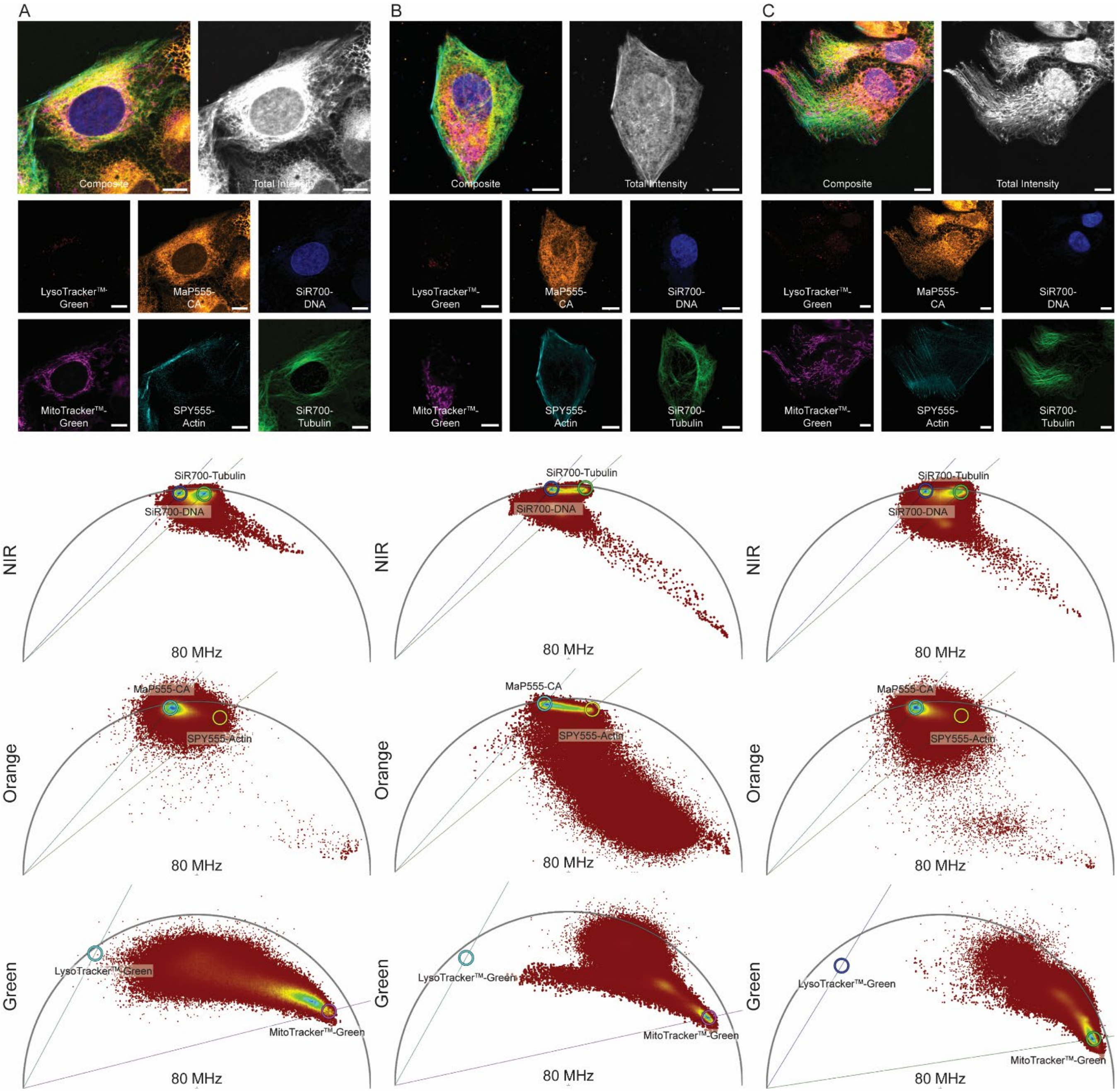

**Supporting Figure S10:** Six species images combining fluorescence lifetime multiplexing in the green, orange and NIR spectral channel. **A-C** U-2 OS cells stably expressing CalR-HaloTag7-SNAP-tag-KDEL were labeled with MitoTracker™-Green, LysoTracker™-Green, MaP555-CA, SPY555-Actin, SiR700-DNA and SiR700-Tubulin and imaged in the three spectral channels giving access to six species images. The composite, the total fluorescence intensity, the six individual separated species as well as the corresponding wavelet-filtered phasor plots used for separation are given. Species separation was achieved using the phasor approach by positioning the cluster-circles on the phasor plot at the position of the pure species. Scale bars, 10 µm.

#### Supporting Tables

**Supporting Table S1:** Structures of all 18 synthetic probes tested. In addition, the vendor/reference and their excitation and emission maximum (as specified by the vendor) are given.

| Name | Vendor/ Reference | Ex/Em [nm] | Structure |
|---|---|---|---|
| **MitoTracker™-Green** | ThermoFisher Scientific | 490/516 | 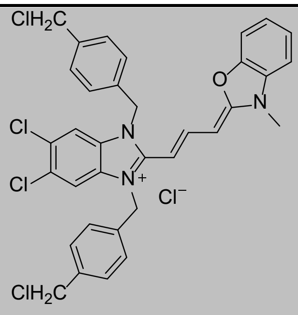 MitoTracker™-Green |
| **LysoTracker™-Green** | ThermoFisher Scientific | 504/511 | 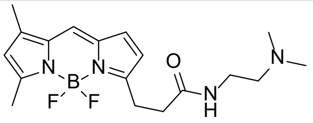 LysoTracker™-Green |
| **SPY555-Actin** | Spirochrome | 555/580 | Unpublished |
| **SPY555-DNA** | Spirochrome | 555/580 | Unpublished |
| **SPY555-Tubulin** | Spirochrome | 555/580 | Unpublished |
| **MaP555-CA** | [1] | 558/578 | 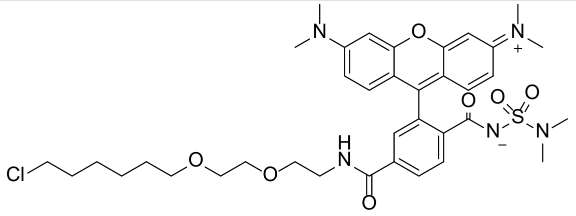 MaP555-CA |
| **MaP555-BG** | [1] | 556/576 | 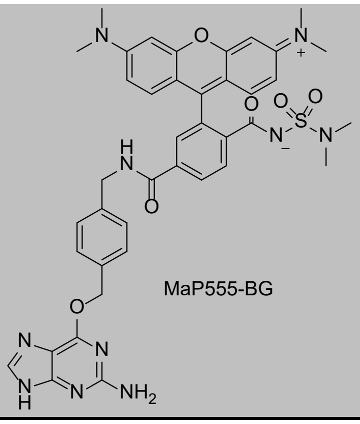 MaP555-BG |

| | | | |
|---|---|---|---|
| **MitoTracker™-Orange** | ThermoFisher Scientific | 554/576 | 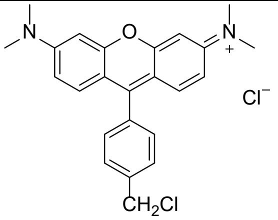 MitoTracker™-Orange |
| **Rhodamine B** | TCI | - | 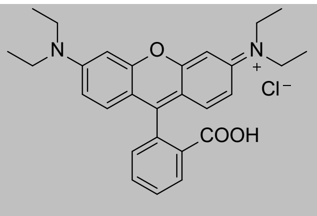 Rhodamine B |
| **BioTracker 560 Orange Lysosomes** | Merck | 532/560 | 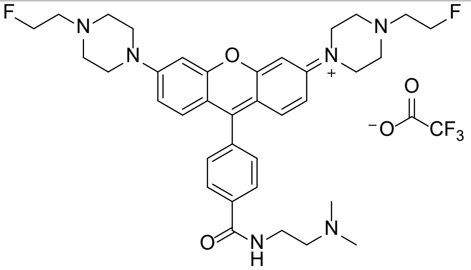 BioTracker 560 Orange Lysosome |
| **MitoTracker™-Red** | ThermoFisher Scientific | 579/599 | 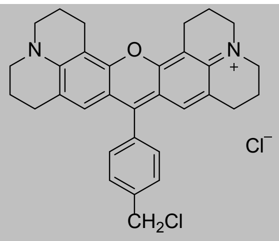 MitoTracker™-Red |
| **LysoTracker™-Red** | ThermoFisher Scientific | 577/590 | 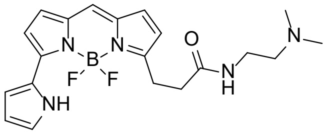 LysoTracker™-Red |
| **SPY620-Actin** | Spirochrome | 619/636 | Unpublished |
| **SPY620-DNA** | Spirochrome | 618/636 | Unpublished |
| **MaP618-CA** | [1] | 618/635 | 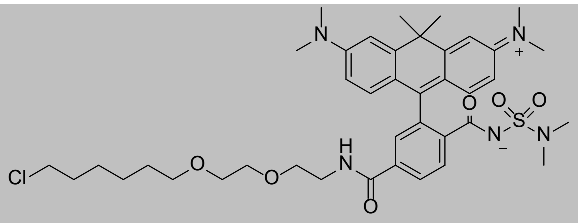 MaP618-CA |

| | | | |
|---|---|---|---|
| **SiR-DNA** | Spirochrome[2] | 652/674 | 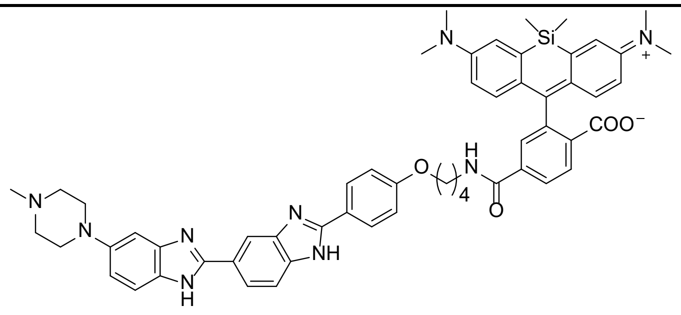 SiR-DNA |
| **SiR-Tubulin** | Spirochrome[3] | 652/674 | 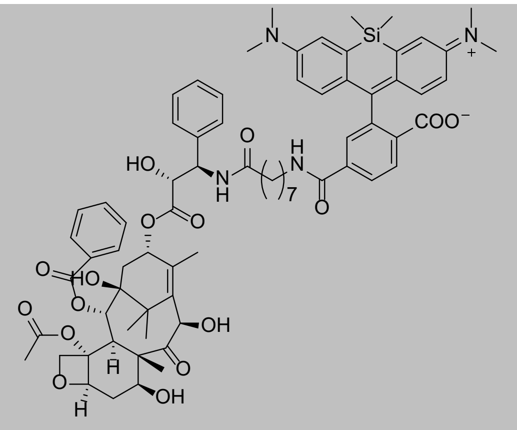 SiR-Tubulin |
| **SiR-Actin** | Spirochrome[3] | 652/674 | 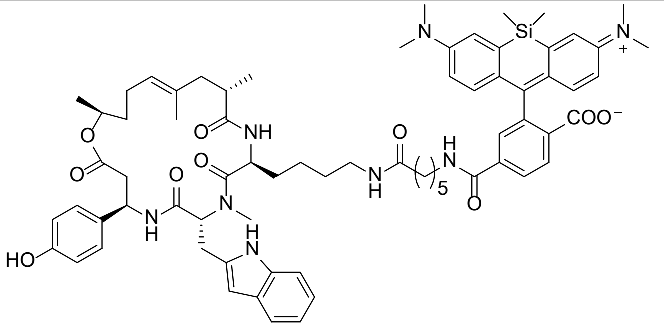 SiR-Actin |
| **SiR700-DNA** | Spirochrome[4] | 689/716 | 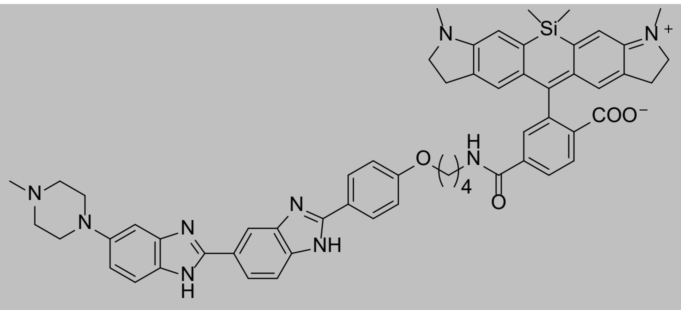 SiR700-DNA |

| | | | |
|---|---|---|---|
| **SiR700-Tubulin** | Spirochrome[4] | 689/716 | 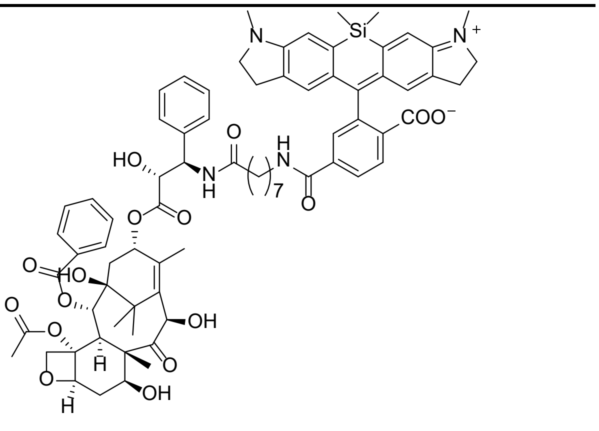 SiR700-Tubulin |
| **SiR700-Actin** | Spirochrome[4] | 689/716 | 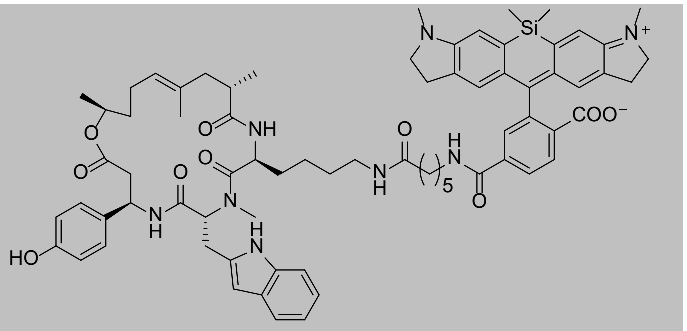 SiR700-Actin |

**Supporting Table S2:** Comparison of intensity weighted fluorescence lifetimes (τ) of different synthetic probes in three different cell lines. Fluorescence lifetimes were measured by FLIM (mean±s.e.m., *N* = 4 field of views (FOVs) from 2 biological replicates, average *N* = 12 FOVs from 6 biological replicates).[2] bi-exponential fit, [3] tri-exponential fit, * tail fit (all others n-exponential reconvolution fit).

| | **Fluorophore** | **U-2 OS [ns]** | | **HeLa [ns]** | | **HEK 293 [ns]** | | **Average [ns]** | |
|---|---|---|---|---|---|---|---|---|---|
| | | τ | s.e.m. | τ | s.e.m. | τ | s.e.m. | τ | s.e.m. |
| **Green** | **MitoTracker™-Green** | 0.96[3*] | 0.03 | 0.88[3*] | 0.03 | 0.96[3*] | 0.02 | 0.93 | 0.02 |
| **Green** | **LysoTracker™-Green** | 3.92[2] | 0.06 | 3.78[2] | 0.05 | 4.24[2] | 0.01 | 3.98 | 0.06 |
| **Orange** | **Rhodamine B** | 2.26[3] | 0.03 | 2.12[3] | 0.01 | 2.30[3] | 0.02 | 2.23 | 0.03 |
| **Orange** | **BioTracker Orange Lysosome** | 2.30[2] | 0.07 | 1.80[3] | 0.13 | 2.04[2] | 0.05 | 2.05 | 0.08 |
| **Orange** | **SPY555-Actin** | 1.79[2] | 0.09 | 1.91[2] | 0.02 | 1.88[2] | 0.03 | 1.86 | 0.03 |
| **Orange** | **SPY555-DNA** | 3.00[2] | 0.02 | 3.01[2] | 0.04 | 3.01[2] | 0.01 | 3.01 | 0.01 |
| **Orange** | **SPY555-Tubulin** | 2.00[2] | 0.01 | 1.99[2] | 0.01 | 2.02[2] | 0.01 | 2.00 | 0.01 |
| **Red** | **LysoTracker™-Red** | 3.88[2] | 0.07 | 3.57[2] | 0.09 | 3.63[2] | 0.06 | 3.70 | 0.06 |
| **Red** | **SPY620-Actin** | 2.93[2] | 0.01 | 2.94[2] | 0.01 | 2.88[2] | 0.03 | 2.91 | 0.01 |
| **Red** | **SPY620-DNA** | 3.62[2] | 0.01 | 3.59[2] | 0.01 | 3.61[2] | 0.01 | 3.60 | 0.01 |
| **NIR** | **SiR700-Actin** | 1.90[2] | 0.01 | 1.88[2] | 0.01 | 1.93[2] | 0.01 | 1.90 | 0.01 |
| **NIR** | **SiR700-DNA** | 2.28[2] | 0.01 | 2.26[2] | 0.01 | 2.30[2] | 0.01 | 2.28 | 0.01 |
| **NIR** | **SiR700-Tubulin** | 1.90[3] | 0.01 | 1.90[3] | 0.01 | 2.02[3] | 0.01 | 1.94 | 0.02 |

**Supporting Table S3:** Comparison of intensity weighted fluorescence lifetimes (τ) of MaP555-BG on SNAPf-tag fused to different proteins of interest. H2B: nucleus, labeling histone 2B, LMNB1: nuclear lamina, labeling Lamin B1, Cytosol: untargeted (no fusion), NES: cytosol, fusion with a nuclear export signal, Tomm20: outer mitochondrial membrane, labeling the membrane receptor Tomm20, COX8: inner mitochondrial membrane, labeling the cytochrome c oxidase subunit 8, CalR-KDEL: endoplasmic reticulum, labeling calreticulin and additionally fused to a KDEL targeting peptide, β-4-Gal-T1: Golgi Apparatus, labeling beta-1,4-galactosidase, LAMP1: lysosomes, labeling lysosome-associated membrane glycoprotein 1, SKL: peroxisomes, targeted via a peroxisomal targeting signal, Lyn11: inner leaflet of the plasma membrane, labeling tyrosine protein kinase Lyn11, Ig-κ-PDGF: outer plasma membrane, labeling platelet-derived growth factor receptor and additionally fused to an immunoglobulin kappa light chain leader sequence, CEP41: microtubules, labeling the microtubule-binding protein CEP41, lifeact: filamentous actin (F-actin), labeling the actin-binding peptide lifeact. Fluorescence lifetime was measured by FLIM (mean±s.e.m., *N* = 4 FOVs from 2 biological replicates). Unless otherwise stated all mono-exponential fits. [2] bi-exponential. Data for HaloTag7 from reference[5].

| Fluorophore | SNAP-tag [ns] | | HaloTag7 [ns] | |
|---|---|---|---|---|
| | τ | s.e.m. | τ | s.e.m. |
| **H2B** | 2.55[2] | 0.03 | 2.41 | 0.01 |
| **LMNB1** | 2.57[2] | 0.04 | 2.38 | 0.00 |
| **Cytosol** | 2.49[2] | 0.01 | 2.33 | 0.01 |
| **NES** | 2.57[2] | 0.02 | 2.41[2] | 0.01 |
| **Tomm20** | 2.51[2] | 0.02 | 2.38 | 0.00 |
| **COX8** | 2.46[2] | 0.04 | 2.40[2] | 0.02 |
| **CalR-KDEL** | 2.52[2] | 0.03 | 2.36 | 0.00 |
| **β4Gal-T1** | 2.58[2] | 0.14 | 2.35 | 0.03 |
| **LAMP1** | 2.62[2] | 0.02 | 2.38 | 0.00 |
| **SKL** | 2.40[2] | 0.05 | 2.37 | 0.01 |
| **Lyn11** | 2.37[2] | 0.05 | 2.37[2] | 0.01 |
| **Ig-κ-PDGF** | 2.34[2] | 0.03 | 2.31[2] | 0.01 |
| **CEP41** | 2.40[2] | 0.06 | 2.51[2] | 0.02 |
| **Lifeact** | 2.48[2] | 0.03 | 2.36[2] | 0.01 |
| **Average** | 2.49[2] | 0.06 | 2.38 | 0.01 |

**Supporting Table S4:** Plasmids used and generated as well as the stable cell line derived thereof.

| Name | Addgene# | Plasmid | Gene | Localization tag Addgene# | Stable cell lines |
|---|---|---|---|---|---|
| **pCDNA5/FRT/TO_H2B-SNAPf-tag** | 181998 | pCDNA5/FRT/TO | H2B-SNAPf-tag | 135444[6] | - |
| **pCDNA5/FRT/TO_SNAPf-tag-LMNB1** | 181999 | pCDNA5/FRT/TO | SNAPf-tag-LMNB1 | 55069 | - |
| **pCDNA5/FRT/TO_SNAPf-tag** | 182000 | pCDNA5/FRT/TO | SNAPf-tag | 167271[7] | - |
| **pCDNA5/FRT/TO_NES-SNAPf-tag** | 182001 | pCDNA5/FRT/TO | NES-SNAPf-tag | 101061[8] | - |
| **pCDNA5/FRT/TO_TOMM20-SNAPf-tag** | 182002 | pCDNA5/FRT/TO | TOMM20-SNAPf-tag | 135443[6] | - |
| **pCDNA5/FRT/TO_COX8-SNAPf-tag** | 182003 | pCDNA5/FRT/TO | COX8-SNAPf-tag | 113916[9] | - |
| **pCDNA5/FRT/TO_CalR-SNAPf-tag -KDEL** | 182004 | pCDNA5/FRT/TO | CalR-SNAPf-tag-KDEL | synthetic | - |
| **pCDNA5/FRT/TO_β4Gal-T1-SNAPf-tag** | 182005 | pCDNA5/FRT/TO | β4Gal-T1-SNAPf-tag | 36205[10] | - |
| **pCDNA5/FRT/TO_LAMP1-SNAPf-tag** | 182006 | pCDNA5/FRT/TO | LAMP1-SNAPf-tag | 34831[11] | - |
| **pCDNA5/FRT/TO_SNAPf-tag-SKL** | 182007 | pCDNA5/FRT/TO | SNAPf-tag-SKL | synthetic | - |
| **pCDNA5/FRT/TO_Lyn11-SNAPf-tag** | 182008 | pCDNA5/FRT/TO | Lyn11-SNAPf-tag | synthetic | - |
| **pCDNA5/FRT/TO_Ig-κ-SNAPf-tag-PDGFR** | 182009 | pCDNA5/FRT/TO | Ig-κ-SNAPf-tag-PDGFR | [12] | - |
| **pCDNA5/FRT/TO_CEP41-SNAPf** | 182010 | pCDNA5/FRT/TO | CEP41-SNAPf | 135446[6] | - |
| **pCDNA5/FRT/TO_Lifeact-SNAPf-tag** | 182011 | pCDNA5/FRT/TO | Lifeact-SNAPf-tag | aynthetic | - |
| **pCDNA5/FRT_CalR-HaloTag7-P30-SNAP-tag-KDEL** | 182012 | pCDNA5/FRT | CalR-SNAPf-tag-KDEL | synthetic | U-2 OS Flp-In TREx |

**Supporting Table S5:** Fluorescence microscopy data acquisition parameters. *See methods for details.

| Image | Probe | Microscope | Excitation [nm] | Pixel dwell time [µs] | Pinhole [Airy Units] | Objective | Pixel size [nm] | Size [pixels] | Emission [nm] | Comment |
|---|---|---|---|---|---|---|---|---|---|---|
| **Fig. 1C** | MitoTracker™-Green | SP8-FALCON | 489 | 3.16 | 1 | 40x1.10 water | 142 | 512x512 | 510-540 | 80 MHz<br>500 photons |
| **Fig. 1C** | LysoTracker™-Green | SP8-FALCON | 489 | 3.16 | 1 | 40x1.10 water | 142 | 512x512 | 510-540 | 80 MHz<br>500 photons |
| **Fig. 1C** | RhodB | SP8-FALCON | 550 | 3.16 | 1 | 40x1.10 water | 142 | 512x512 | 570-600 | 80 MHz<br>500 photons |
| **Fig. 1C** | BioTracker Orange Lysosome | SP8-FALCON | 550 | 3.16 | 1 | 40x1.10 water | 142 | 512x512 | 570-600 | 80 MHz<br>500 photons |
| **Fig. 1C** | SPY555-Actin | SP8-FALCON | 550 | 3.16 | 1 | 40x1.10 water | 142 | 512x512 | 570-600 | 80 MHz<br>500 photons |
| **Fig. 1C** | SPY555-DNA | SP8-FALCON | 550 | 3.16 | 1 | 40x1.10 water | 142 | 512x512 | 570-600 | 80 MHz<br>500 photons |
| **Fig. 1C** | SPY555-Tubulin | SP8-FALCON | 550 | 3.16 | 1 | 40x1.10 water | 142 | 512x512 | 570-600 | 80 MHz<br>500 photons |
| **Fig. 1C** | SPY620-Actin | SP8-FALCON | 615 | 3.16 | 1 | 40x1.10 water | 142 | 512x512 | 635-700 | 80 MHz<br>500 photons |
| **Fig. 1C** | SPY620-DNA | SP8-FALCON | 615 | 3.16 | 1 | 40x1.10 water | 142 | 512x512 | 635-700 | 80 MHz<br>500 photons |
| **Fig. 1C** | SiR700-Actin | SP8-FALCON | 670 | 3.16 | 1 | 40x1.10 water | 142 | 512x512 | 710-760 | 80 MHz<br>500 photons |
| **Fig. 1C** | SiR700-DNA | SP8-FALCON | 670 | 3.16 | 1 | 40x1.10 water | 142 | 512x512 | 710-760 | 80 MHz<br>500 photons |
| **Fig. 1C** | SiR700-Tubulin | SP8-FALCON | 670 | 3.16 | 1 | 40x1.10 water | 142 | 512x512 | 710-760 | 80 MHz<br>500 photons |
| **Fig2A** | MitoTracker™-Green<br>LysoTracker™-Green | SP8-FALCON | 489 | 2.30 | 1 | 40x1.10 water | 90 | 704x704 | 510-540 | 80 MHz<br>6 line accumu. |
| **Fig2B** | RhodB<br>SPY555-DNA | SP8-FALCON | 550 | 2.70 | 1 | 40x1.10 water | 102 | 600x600 | 570-600 | 80 MHz<br>10 line accumu. |
| **Fig2C** | SPY620-DNA | SP8-FALCON | 615 | 4.06 | 1 | 40x1.10 water | 112 | 400x400 | 635-700 | 80 MHz |

| | | | | | | | | | | |
|---|---|---|---|---|---|---|---|---|---|---|
| | SPY620-Actin | | | | | | | | | 10 line accumu. |
| **Fig2D** | SiR700-DNA<br>SiR700-Tubulin | SP8-FALCON | 670 | 8.49 | 1 | 40x1.10 water | 122 | 464x464 | 710-760 | 80 MHz<br>10 line accumu. |
| **Fig3B** | MitoTracker$^{TM}$-Green<br>LysoTracker$^{TM}$-Green | SP8-FALCON | 489 | 2.00 | 1 | 40x1.10 water | 90 | 1160x1160 | 510-540 | 80 MHz<br>10 line accumu. |
| **Fig3B** | CalR-Halo-KDEL<br>MaP555-CA<br>SPY555-Actin | SP8-FALCON | 550 | 2.00 | 1 | 40x1.10 water | 90 | 1160x1160 | 570-600 | 80 MHz<br>10 line accumu. |
| **Fig3B** | SiR700-DNA<br>SiR700-Tubulin | SP8-FALCON | 670 | 2.00 | 1 | 40x1.10 water | 90 | 1160x1160 | 710-760 | 80 MHz<br>10 line accumu. |
| **S1A** | MitoTracker$^{TM}$-Green | SP8-FALCON | 489 | 3.16 | 1 | 40x1.10 water | 142 | 512x512 | 510-540 | 80 MHz<br>500 photons |
| **S1A** | LysoTracker$^{TM}$-Green | SP8-FALCON | 489 | 3.16 | 1 | 40x1.10 water | 142 | 512x512 | 510-540 | 80 MHz<br>500 photons |
| **S1B** | MitoTracker$^{TM}$-Red | SP8-FALCON | 575 | 3.16 | 1 | 40x1.10 water | 142 | 512x512 | 595-625 | 80 MHz<br>500 photons |
| **S1B** | LysoTracker$^{TM}$-Red | SP8-FALCON | 575 | 3.16 | 1 | 40x1.10 water | 142 | 512x512 | 595-625 | 80 MHz<br>500 photons |
| **S1B** | SPY620-Actin | SP8-FALCON | 615 | 3.16 | 1 | 40x1.10 water | 142 | 512x512 | 635-700 | 80 MHz<br>500 photons |
| **S1B** | SPY620-DNA | SP8-FALCON | 615 | 3.16 | 1 | 40x1.10 water | 142 | 512x512 | 635-700 | 80 MHz<br>500 photons |
| **S1C** | RhodB | SP8-FALCON | 550 | 3.16 | 1 | 40x1.10 water | 142 | 512x512 | 570-600 | 80 MHz<br>500 photons |
| **S1C** | BioTracker Orange<br>Lysosome | SP8-FALCON | 550 | 3.16 | 1 | 40x1.10 water | 142 | 512x512 | 570-600 | 80 MHz<br>500 photons |
| **S1C** | SPY555-Actin | SP8-FALCON | 550 | 3.16 | 1 | 40x1.10 water | 142 | 512x512 | 570-600 | 80 MHz<br>500 photons |
| **S1C** | SPY555-DNA | SP8-FALCON | 550 | 3.16 | 1 | 40x1.10 water | 142 | 512x512 | 570-600 | 80 MHz<br>500 photons |
| **S1C** | SPY555-Tubulin | SP8-FALCON | 550 | 3.16 | 1 | 40x1.10 water | 142 | 512x512 | 570-600 | 80 MHz<br>500 photons |
| **S1C** | MitoTracker$^{TM}$-<br>Orange | SP8-FALCON | 550 | 3.16 | 1 | 40x1.10 water | 142 | 512x512 | 570-600 | 80 MHz<br>500 photons |
| **S1D** | SiR-DNA | SP8-FALCON | 631 | 3.16 | 1 | 40x1.10 water | 569 | 512x512 | 650-700 | 80 MHz<br>1,000<br>4 line accumu. |

| | | | | | | | | | | |
|---|---|---|---|---|---|---|---|---|---|---|
| **S1D** | SiR-Actin | SP8-FALCON | 631 | 3.16 | 1 | 40x1.10 water | 569 | 512x512 | 650-700 | 80 MHz<br>1,000<br>4 line accumu. |
| **S1D** | SiR-Tubulin | SP8-FALCON | 631 | 3.16 | 1 | 40x1.10 water | 569 | 512x512 | 650-700 | 80 MHz<br>1,000<br>4 line accumu. |
| **S1E** | SiR700-Actin | SP8-FALCON | 670 | 3.16 | 1 | 40x1.10 water | 142 | 512x512 | 710-760 | 80 MHz<br>500 photons |
| **S1E** | SiR700-DNA | SP8-FALCON | 670 | 3.16 | 1 | 40x1.10 water | 142 | 512x512 | 710-760 | 80 MHz<br>500 photons |
| **S1E** | SiR700-Tubulin | SP8-FALCON | 670 | 3.16 | 1 | 40x1.10 water | 142 | 512x512 | 710-760 | 80 MHz<br>500 photons |
| **S2A** | MitoTracker™-Green or<br>LysoTracker™-Green | SP8 | * | 3.16 | 1 | 40x1.10 water | 142 | 512x512 | * | Spectra |
| **S2B** | MaP618-CA or<br>SPY620-DNA or<br>SPY620-Actin | SP8 | * | 3.16 | 1 | 40x1.10 water | 142 | 512x512 | * | Spectra |
| **S2B** | LysoTracker™-Red | SP8 | * | 3.16 | 1 | 40x1.10 water | 142 | 512x512 | * | Spectra |
| **S2C-D** | MaP555-CA or<br>MaP555-BG or<br>SPY555-DNA or<br>SPY555-Actin or<br>SPY555-Tubulin or<br>Rhodamine B or<br>Bio Tracker Orange Lysosome | SP8 | * | 3.16 | 1 | 40x1.10 water | 142 | 512x512 | * | Spectra |
| **S2E** | SiR700-DNA or<br>SiR700-Actin or<br>SiR700-Tubulin | SP8 | * | 3.16 | 1 | 40x1.10 water | 142 | 512x512 | * | Spectra |
| **S3A** | MitoTracker™-Green<br>LysoTracker™-Green | SP8-FALCON | 489 | 2.30 | 1 | 40x1.10 water | 90 | 704x704 | 510-540 | 80 MHz<br>6 line accumu. |
| **S3B** | SPY620-DNA<br>SPY620-Actin | SP8-FALCON | 615 | 4.06 | 1 | 40x1.10 water | 112 | 400x400 | 635-700 | 80 MHz<br>10 line accumu. |
| **S3C** | SiR700-DNA<br>SiR700-Actin | SP8-FALCON | 670 | 6.65 | 1 | 40x1.10 water | 123 | 592x592 | 710-760 | 80 MHz<br>10 line accumu. |
| **S3D** | SiR700-DNA<br>SiR700-Tubulin | SP8-FALCON | 670 | 8.49 | 1 | 40x1.10 water | 122 | 464x464 | 710-760 | 80 MHz<br>10 line accumu. |

| | | | | | | | | | | |
|---|---|---|---|---|---|---|---|---|---|---|
| **S4A** | Bio Tracker Orange Lysosome<br>SPY555-DNA | SP8-FALCON | 550 | 2.85 | 1 | 40x1.10 water | 101 | 568x568 | 570-600 | 80 MHz<br>10 line accumu. |
| **S4B** | SPY555-Actin<br>SPY555-DNA | SP8-FALCON | 550 | 2.50 | 1 | 40x1.10 water | 101 | 648x648 | 570-600 | 80 MHz<br>10 line accumu. |
| **S4C** | RhodB<br>SPY555-DNA | SP8-FALCON | 550 | 2.70 | 1 | 40x1.10 water | 102 | 600x600 | 570-600 | 80 MHz<br>10 line accumu. |
| **S4D** | SPY555-Tubulin<br>SPY555-DNA | SP8-FALCON | 550 | 1.56 | 1 | 40x1.10 water | 101 | 1040x1040 | 570-600 | 80 MHz<br>10 line accumu. |
| **S5A** | Bio Tracker Orange Lysosome<br>SPY555-DNA | SP8-FALCON | 550 | 3.69 | 1 | 40x1.10 water | 101 | 440x440 | 570-600 | 80 MHz<br>10 line accumu. |
| **S5B** | SPY555-Tubulin<br>SPY555-DNA | SP8-FALCON | 550 | 9.85 | 1 | 40x1.10 water | 100 | 400x400 | 570-600 | 80 MHz<br>10 line accumu. |
| **S5C** | Bio Tracker Orange Lysosome<br>SPY555-DNA | SP8-FALCON | 550 | 4.06 | 1 | 40x1.10 water | 101 | 400x400 | 570-600 | 80 MHz<br>10 line accumu. |
| **S5D** | SPY555-Tubulin<br>SPY555-DNA | SP8-FALCON | 550 | 7.24 | 1 | 40x1.10 water | 100 | 544x544 | 570-600 | 80 MHz<br>10 line accumu. |
| **S6A** | SPY555-DNA<br>CalR-Halo-KDEL<br>MaP555-CA | SP8-FALCON | 550 | 2.25 | 1 | 40x1.10 water | 101 | 728x728 | 570-600 | 80 MHz<br>10 line accumu. |
| **S6B** | SPY555-Actin<br>CalR-Halo-KDEL<br>MaP555-CA | SP8-FALCON | 550 | 1.70 | 1 | 40x1.10 water | 101 | 952x952 | 570-600 | 80 MHz<br>10 line accumu. |
| **S6C** | SPY555-DNA<br>CalR-SNAP-KDEL<br>MaP555-BG | SP8-FALCON | 550 | 2.25 | 1 | 40x1.10 water | 101 | 720x720 | 570-600 | 80 MHz<br>10 line accumu. |
| **S6D** | SPY555-Actin<br>CalR-SNAP-KDEL<br>MaP555-BG | SP8-FALCON | 550 | 1.93 | 1 | 40x1.10 water | 101 | 840x840 | 570-600 | 80 MHz<br>10 line accumu. |
| **S7** | SPY620-DNA<br>CalR-Halo-KDEL<br>MaP618-CA | SP8-FALCON | 615 | 2.50 | 1 | 40x1.10 water | 114 | 648x648 | 635-700 | 80 MHz<br>10 line accumu. |
| **S8A** | MitoTracker™-Green<br>LysoTracker™-Green | SP8-FALCON | 489 | 1.96 | 1 | 40x1.10 water | 90 | 824x824 | 510-540 | 80 MHz<br>10 line accumu. |
| **S8A** | SPY555-DNA<br>SPY555-Tubulin | SP8-FALCON | 550 | 1.96 | | 40x1.10 water | 90 | 824x824 | 570-600 | 80 MHz<br>10 line accumu. |

| | | | | | | | | | | |
|---|---|---|---|---|---|---|---|---|---|---|
| **S8B** | MitoTracker™-Green<br>LysoTracker™-Green | SP8-FALCON | 489 | 1.83 | 1 | 40x1.10 water | 90 | 888x888 | 510-540 | 80 MHz<br>10 line accumu. |
| **S8B** | SPY555-DNA<br>SPY555-Actin | SP8-FALCON | 550 | 1.83 | 1 | 40x1.10 water | 90 | 888x888 | 570-600 | 80 MHz<br>10 line accumu. |
| **S9** | MitoTracker™-Green<br>LysoTracker™-Green | SP8-FALCON | 489 | 1.89 | 1 | 40x1.10 water | 90 | 856x856 | 510-540 | 80 MHz<br>12 line accumu. |
| **S9** | SPY620-DNA<br>SPY620-Actin | SP8-FALCON | 615 | 1.89 | 1 | 40x1.10 water | 90 | 856x856 | 635-700 | 80 MHz<br>12 line accumu. |
| **S10A** | MitoTracker™-Green<br>LysoTracker™-Green | SP8-FALCON | 489 | 2.01 | 1 | 40x1.10 water | 90 | 776x776 | 510-540 | 80 MHz<br>10 line accumu. |
| **S10A** | CalR-Halo-KDEL<br>MaP555-CA<br>SPY555-Actin | SP8-FALCON | 550 | 2.01 | 1 | 40x1.10 water | 90 | 776x776 | 570-600 | 80 MHz<br>10 line accumu. |
| **S10A** | SiR700-DNA<br>SiR700-Tubulin | SP8-FALCON | 670 | 2.01 | 1 | 40x1.10 water | 90 | 776x776 | 710-760 | 80 MHz<br>10 line accumu. |
| **S10B** | MitoTracker™-Green<br>LysoTracker™-Green | SP8-FALCON | 489 | 1.46 | 1 | 40x1.10 water | 90 | 712x712 | 510-540 | 80 MHz<br>10 line accumu. |
| **S10B** | CalR-Halo-KDEL<br>MaP555-CA<br>SPY555-Actin | SP8-FALCON | 550 | 1.46 | 1 | 40x1.10 water | 90 | 712x712 | 570-600 | 80 MHz<br>10 line accumu. |
| **S10B** | SiR700-DNA<br>SiR700-Tubulin | SP8-FALCON | 670 | 1.46 | 1 | 40x1.10 water | 90 | 712x712 | 710-760 | 80 MHz<br>10 line accumu. |
| **S10C** | MitoTracker™-Green<br>LysoTracker™-Green | SP8-FALCON | 489 | 2.00 | 1 | 40x1.10 water | 90 | 1160x1160 | 510-540 | 80 MHz<br>10 line accumu. |
| **S10C** | CalR-Halo-KDEL<br>MaP555-CA<br>SPY555-Actin | SP8-FALCON | 550 | 2.00 | 1 | 40x1.10 water | 90 | 1160x1160 | 570-600 | 80 MHz<br>10 line accumu. |
| **S10C** | SiR700-DNA<br>SiR700-Tubulin | SP8-FALCON | 670 | 2.00 | 1 | 40x1.10 water | 90 | 1160x1160 | 710-760 | 80 MHz<br>10 line accumu. |
| **Tab S2** | MitoTracker™-Green | SP8-FALCON | 489 | 3.16 | 1 | 40x1.10 water | 142 | 512x512 | 510-540 | 80 MHz<br>500 photons |
| **Tab S2** | LysoTracker™-Green | SP8-FALCON | 489 | 3.16 | 1 | 40x1.10 water | 142 | 512x512 | 510-540 | 80 MHz<br>500 photons |
| **Tab S2** | RhodB | SP8-FALCON | 550 | 3.16 | 1 | 40x1.10 water | 142 | 512x512 | 570-600 | 80 MHz<br>500 photons |
| **Tab S2** | BioTracker Orange<br>Lysosome | SP8-FALCON | 550 | 3.16 | 1 | 40x1.10 water | 142 | 512x512 | 570-600 | 80 MHz<br>500 photons |
| **Tab S2** | SPY555-Actin | SP8-FALCON | 550 | 3.16 | 1 | 40x1.10 water | 142 | 512x512 | 570-600 | 80 MHz |

| | | | | | | | | | | |
|---|---|---|---|---|---|---|---|---|---|---|
| | | | | | | | | | | 500 photons |
| **Tab S2** | SPY555-DNA | SP8-FALCON | 550 | 3.16 | 1 | 40x1.10 water | 142 | 512x512 | 570-600 | 80 MHz<br>500 photons |
| **Tab S2** | SPY555-Tubulin | SP8-FALCON | 550 | 3.16 | 1 | 40x1.10 water | 142 | 512x512 | 570-600 | 80 MHz<br>500 photons |
| **Tab S2** | LysoTracker$^{TM}$-Red | SP8-FALCON | 575 | 3.16 | 1 | 40x1.10 water | 142 | 512x512 | 595-625 | 80 MHz<br>500 photons |
| **Tab S2** | SPY620-Actin | SP8-FALCON | 615 | 3.16 | 1 | 40x1.10 water | 142 | 512x512 | 635-700 | 80 MHz<br>500 photons |
| **Tab S2** | SPY620-DNA | SP8-FALCON | 615 | 3.16 | 1 | 40x1.10 water | 142 | 512x512 | 635-700 | 80 MHz<br>500 photons |
| **Tab S2** | SiR700-Actin | SP8-FALCON | 670 | 3.16 | 1 | 40x1.10 water | 142 | 512x512 | 710-760 | 80 MHz<br>500 photons |
| **Tab S2** | SiR700-DNA | SP8-FALCON | 670 | 3.16 | 1 | 40x1.10 water | 142 | 512x512 | 710-760 | 80 MHz<br>500 photons |
| **Tab S2** | SiR700-Tubulin | SP8-FALCON | 670 | 3.16 | 1 | 40x1.10 water | 142 | 512x512 | 710-760 | 80 MHz<br>500 photons |
| **Tab S3** | H2B-SNAP<br>MaP555-BG | SP8-FALCON | 550 | 3.16 | 1 | 40x1.10 water | 142 | 512x512 | 570-600 | 80 MHz<br>500 photons |
| **Tab S3** | SNAP-LMNB1<br>MaP555-BG | SP8-FALCON | 550 | 5.64 | 1 | 40x1.10 water | 101 | 288x288 | 570-600 | 80 MHz<br>500 photons |
| **Tab S3** | SNAP<br>MaP555-BG | SP8-FALCON | 550 | 3.16 | 1 | 40x1.10 water | 142 | 512x512 | 570-600 | 80 MHz<br>500 photons |
| **Tab S3** | NES-SNAP<br>MaP555-BG | SP8-FALCON | 550 | 3.16 | 1 | 40x1.10 water | 142 | 512x512 | 570-600 | 80 MHz<br>500 photons |
| **Tab S3** | Tomm20-SNAP<br>MaP555-BG | SP8-FALCON | 550 | 3.16 | 1 | 40x1.10 water | 142 | 512x512 | 570-600 | 80 MHz<br>500 photons |
| **Tab S3** | COX8-SNAP<br>MaP555-BG | SP8-FALCON | 550 | 3.16 | 1 | 40x1.10 water | 142 | 512x512 | 570-600 | 80 MHz<br>500 photons |
| **Tab S3** | CalR-SNAP-KDEL<br>MaP555-BG | SP8-FALCON | 550 | 3.16 | 1 | 40x1.10 water | 142 | 512x512 | 570-600 | 80 MHz<br>500 photons |
| **Tab S3** | β4Gal-T1-SNAP<br>MaP555-BG | SP8-FALCON | 550 | 3.16 | 1 | 40x1.10 water | 142 | 512x512 | 570-600 | 80 MHz<br>500 photons |
| **Tab S3** | LAMP1-SNAP<br>MaP555-BG | SP8-FALCON | 550 | 3.16 | 1 | 40x1.10 water | 142 | 512x512 | 570-600 | 80 MHz<br>500 photons |
| **Tab S3** | SNAP-SKL<br>MaP555-BG | SP8-FALCON | 550 | 7.69 | 1 | 40x1.10 water | 142 | 512x512 | 570-600 | 80 MHz<br>500 photons |
| **Tab S3** | Lyn11-SNAP | SP8-FALCON | 550 | 3.16 | 1 | 40x1.10 water | 142 | 512x512 | 570-600 | 80 MHz |

| | | | | | | | | | | |
|---|---|---|---|---|---|---|---|---|---|---|
| | MaP555-BG | | | | | | | | | 500 photons |
| **Tab S3** | Ig-κ-SNAP-PDGFR MaP555-BG | SP8-FALCON | 550 | 3.16 | 1 | 40x1.10 water | 142 | 512x512 | 570-600 | 80 MHz 500 photons |
| **Tab S3** | CEP41-SNAP MaP555-BG | SP8-FALCON | 550 | 3.16 | 1 | 40x1.10 water | 142 | 512x512 | 570-600 | 80 MHz 500 photons |
| **Tab S3** | Lifeact-SNAP MaP555-BG | SP8-FALCON | 550 | 3.16 | 1 | 40x1.10 water | 142 | 512x512 | 570-600 | 80 MHz 500 photons |
